# JAK-STAT Pathway Heterogeneity Governs Immunotherapy Response in Breast Cancer

**DOI:** 10.64898/2026.02.03.703506

**Authors:** Jianbo Zhou, Heng Zhang, Hailin Tang, Lei Yu, Fu Peng

## Abstract

The JAK-STAT pathway (JSP) is a well-known oncogenic cascade; however, recent clinical trials have detected JSP upregulation in breast cancer following anti-PD1 immunotherapy. This paradoxical observation warrants further investigation into JSP’s intercellular heterogeneity, tumor dynamics, molecular mechanisms, and clinical implications for immunotherapy. JSP expression showed dynamic shifts during breast cancer progression, with higher levels in T cells and para-cancerous epithelial cells. In tumor cells, elevated JSP highly correlated with malignant phenotypes. JSP-high tumor cells overexpressed oncogenic pathways, while exhibiting increased immunosuppressive signaling via MIF-CD74 signaling axis. In T cells, higher JSP levels were associated with enhanced cytotoxic activity, improved differentiation, and reduced exhaustion, reflecting robust anti-tumor immunity. Analysis of immunotherapy datasets revealed that higher JSP levels were associated with improved responses towards PD-1 inhibitors, particularly in triple-negative breast cancer (TNBC) patients, with JSP serving as a predictive biomarker for immunotherapy sensitivity. As a key JSP component, STAT4 exerts dual roles in breast cancer: it drives tumorigenesis in malignant cells, sustains breast epithelial cell proliferation, and bolsters T cell anti-tumor functionality--while also acting as a highly accurate biomarker for predicting immunotherapy response. This indicates that JSP targeting demands a nuanced approach: broad inhibition may impair anti-tumor immunity, and optimized therapeutic strategies paired with precise biomarkers are critical to maximize JSP’s utility in breast cancer immunotherapy. Our findings highlight JSP’s functional heterogeneity in epithelial, tumor, and T cells, with high JSP activity correlating with enhanced immunotherapy efficacy in breast cancer.

**Graphic Abstract:** 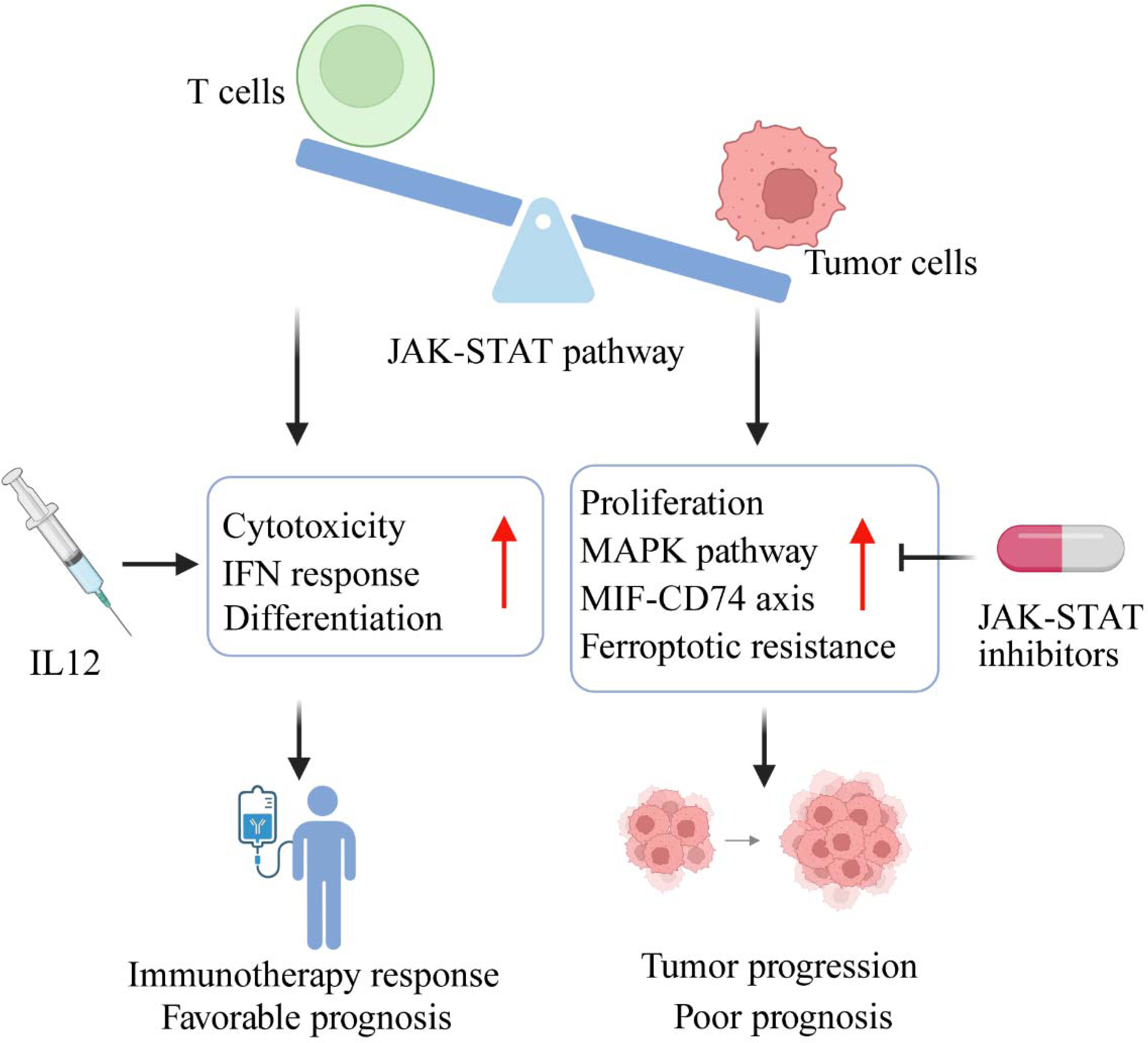

## 1. Introduction

Breast cancer is the most common cancer in women, accounting for 32% (321,910 cases) of new cancer diagnoses and 14% (42,140 cases) of cancer-related deaths among women in United States, 2026 ^1^. Globally, approximately 2.3 million new breast cancer cases and 670,000 breast cancer-related deaths are reported in 2022 ^2^. Breast cancer is commonly classified by immunohistochemistry (IHC) based on the expression of estrogen receptor (ER), progesterone receptor (PR), and human epidermal growth factor receptor 2 (HER2). Using these markers, tumors are typically divided into four major subtypes—luminal A, luminal B, HER2□positive, and triple□negative breast cancer (TNBC). Clinically, this scheme is often simplified into three groups: Luminal (ER and/or PR positive, HER2 negative), HER2□positive, and triple□negative breast cancer (ER, PR and HER2 all negative; TNBC) ^3^. TNBC accounts for approximately 15% of all breast cancers, with a rising overall incidence in recent years and higher prevalence among young women ^2^. It can be further classified into distinct molecular subgroups, including basal□like (BL1 and BL2), claudin□low, mesenchymal (MES), luminal androgen receptor (LAR), and immunomodulatory (IM) types ^4,5^. Among these, basal□like subtypes predominate, with BL1 and BL2 representing the largest proportions. An alternative molecular classification based on genomic biomarkers divides TNBC into four subtypes: LAR, IM, basal□like immune□suppressed (BLIS), and mesenchymal□like (MES) ^6^. TNBC is characterized by aggressive behavior, a high risk of early relapse, and a propensity to present at advanced stages. Histologically, TNBCs are often poorly differentiated, highly proliferative, and heterogeneous, encompassing subsets with variable prognoses. Immunohistochemically, TNBC can be categorized as basal or non□basal: basal tumors typically express cytokeratins CK5/6 and EGFR, whereas non□basal tumors do not express CK5/6. Notably, approximately 80% of tumors associated with BRCA1/BRCA2 alterations fall into the TNBC category ^7^. Current standard treatments include surgery, radiotherapy, neoadjuvant chemotherapy or / and immunotherapy with immune checkpoint inhibitors (ICI) (e.g., anti-PD-1: Pembrolizumab; anti-PD-L1: Atezolizumab; anti-CTLA4: Tremelimumab) ^8,9^. Although immunotherapy has reshaped breast cancer treatment patterns and improved patient’s survival benefits, heterogeneity in treatment responses and the lack of robust predictive biomarkers continue to limit its optimal clinical application ^10–12^.

The JAK-STAT (Janus Kinase-Signal Transducer and Activator of Transcription) pathway is composed of cytokine receptors (e.g., IL-2, IL-6, IL-10, PDGF, EGF) ^13^, the JAK family (JAK1, JAK2, JAK3, and TYK2), the STAT family (STAT1-6, STAT5a and STAT5b), and negative regulatory factors. These negative regulators, including the SOCS family, PIAS family, and protein tyrosine phosphatases (PTPs), inhibit the activation of the JAK-STAT pathway by suppressing the phosphorylation of JAK or / and STAT proteins ^14^, while PP2A and NPM-ALK maintain STAT3 phosphorylation ^15^. In the canonical working model of JAK-STAT signaling, cytokine binding to its cognate receptor (e.g., gp130, EpoR, TNF-R1, IL-17R, IL-10R) triggers the phosphorylation and activation of JAKs. Activated JAKs then phosphorylate STATs, inducing their dimerization. The dimeric STATs translocate into the nucleus and regulate the transcription of target genes ^13,16^. This pathway is regarded as one of the major abnormally activated oncogenic pathways in breast cancer, contributing to tumor proliferation, invasion, and metastasis ^8^ and collaborating with TGF-β, MAPK, Nocth, PI3K/AKT/mTOR and NF-κB signaling pathway ^13^.

Interestingly, the JAK-STAT pathway operates as a double□edged sword in tumor immunity, mediating both pro□ and anti□tumor programs within the tumor microenvironment (TME) ^17^. On the pro□tumor side, signalling through STAT3 and STAT5 drives immunosuppressive networks in helper T cells, regulatory T cells and myeloid□derived suppressor cells (MDSCs), with downstream cytokines such as IL□6 and IL□23 reinforcing tumor□promoting inflammation and immune evasion. Conversely, anti□tumor activity in the TME is largely orchestrated by STAT1 and STAT4. Th1□associated axes — notably IL□12/STAT4/IL□2 and IFN□γ/STAT1/IFN□γ — foster expansion and activation of CD8^+^ T cells and NK cells. In above cytotoxic lymphocytes, IL□12-STAT4 and IFN□α/β-STAT1 signalling induce granzyme B and IFN□γ production, initiating tumor cell killing ^18^. In dendritic cells (DCs), type I interferon signalling via STAT1 (and in some contexts STAT3) upregulates key effector mediators such as IL□12, TNF□α and interferons, thereby enhancing antigen presentation and T□cell priming ^17^. Additionally, STAT3 maintains proliferation of naïve CD8^+^ T cells, contributes to the initial expansion of activated CD8^+^ T cells, and facilitates the differentiation from memory CD8^+^ T cells to effectors CD8^+^ T cells through up-regulating the key transcriptional factor Bcl-6 ^19^.

Recent studies have observed activation of JAK-STAT pathway following neoadjuvant immunotherapy (PD-1 inhibitor), and this activation was considered to contribute to treatment response and pathological complete response (pCR) in breast cancer ^20,21^. The apparent paradox — that JAK-STAT signalling can drive tumorigenesis yet becomes hyperactivated following immunotherapy — prompted us to dissect its context□dependent roles within TME. We combined bulk RNA□seq with single□cell RNA□seq from primary tumors and immunotherapy cohorts to map JAK-STAT dynamics during the transition from normal to malignant epithelium and to chart its heterogeneity across epithelial cells and T cells (Fig. 1A). We find that JAK-STAT signalling is essential for epithelial differentiation and T□cell cytotoxicity; in immunotherapy-treated cohorts, the pathway activation is associated with enhanced T□cell-mediated cytotoxicity, and predicts clinical response and T□cell expansion. Multi□omics profiling and experiments further identify STAT4 as a salient marker and mediator of JAK-STAT activity. Altogether, these results delineate distinct biological roles and molecular dependencies of the JAK-STAT axis in tumor microenvironment and argue that precise modulation of this pathway could augment immunotherapy efficacy while limiting harm to normal epithelial cells and T cells.

**Figure 1.**
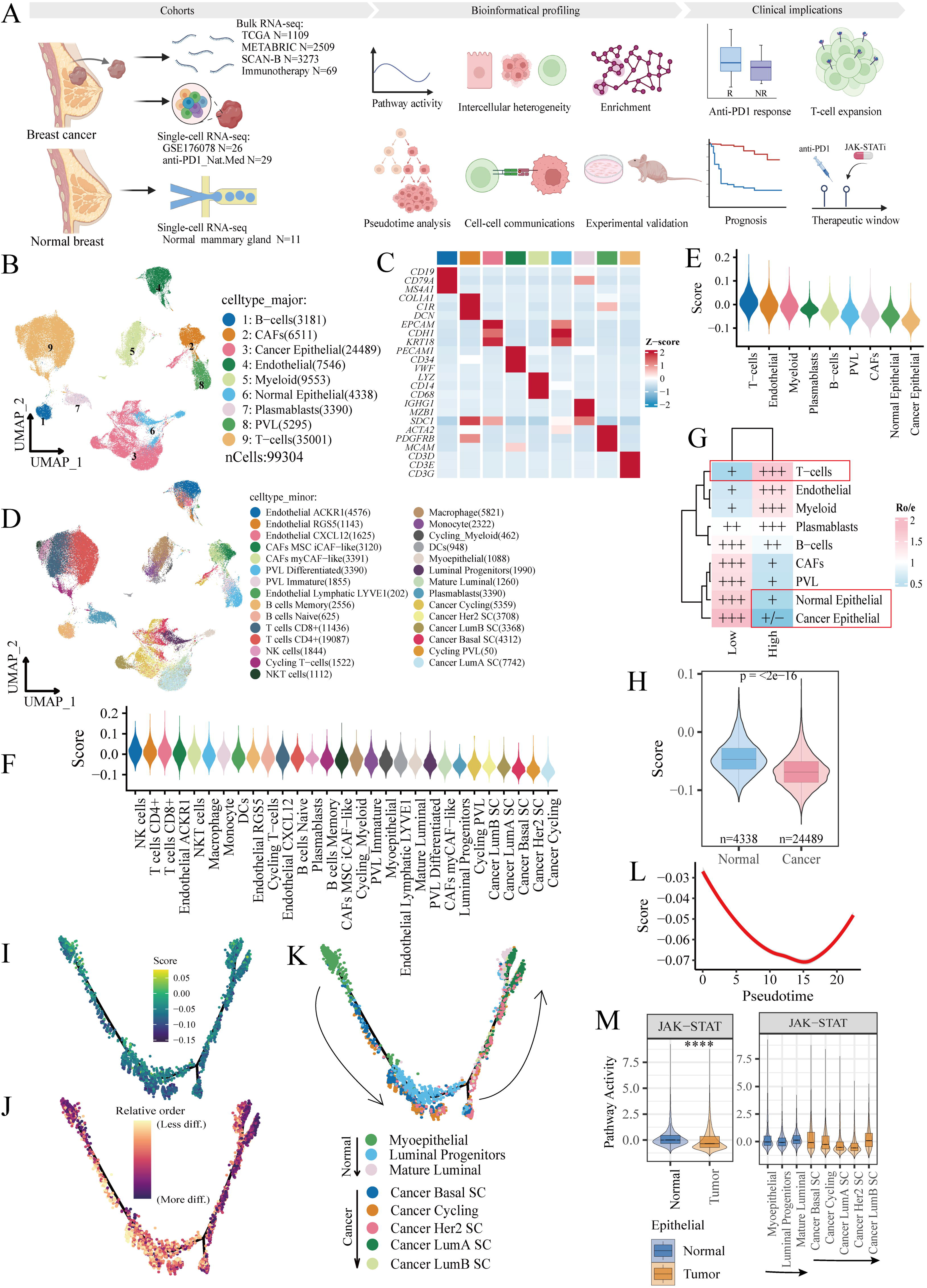
Heterogeneous expression pattern of JAK−STAT pathway in breast cancer. (A) Schematic illustration of the overall experimental and analytical workflow. (B) UMAP plot illustrating the major cell populations derived from the single-cell dataset GSE176078. (C) Core marker gene expressions of major cell populations. (D) UMAP plot depicting the minor cell subpopulations. (E-F) Distribution of JAK-STAT pathway scores across major and minor cell types. (G) Ro/e index for JAK-STAT pathway score. (H) JAK-STAT pathway scores in paracancerous epithelial cells versus cancerous epithelial cells. (I) Pseudotime trajectory analysis of epithelial cell states using Monocle2. (J-K) Pseudotime scores computed by Monocle2 and CytoTRACE2-derived cytotrace scores (in which higher scores indicate lower levels of differentiation). (K) Differentiation trajectories of epithelial cell subtypes. (L) Pseudotime dynamics of the JAK-STAT pathway scores. (M) Activity of the JAK-STAT pathway within epithelial cells.

## 2. Methods and materials

### 2.1. Bioinformatical profiling

#### 2.1.1 Data collection

The annotated human breast cancer single-cell RNA-sequencing dataset (GSE176078) with 26 cases (11 ER^+^, 5 HER2^+^, and 10 TNBC) was downloaded in NCBI’s Gene Expression Omnibus: NCBI-GEO (GSE176078) and analyzed according to previous studies ^22,23^. Human normal breast epithelial single-cell RNA-sequencing dataset was deposited in the NCBI-GEO database (GSE161529), and the .rds format file (SeuratObject_NormEpi.rds) was downloaded in the Figureshare (https://doi.org/10.6084/m9.figshare.17058077) ^24,25^. The single-cell dataset of anti-PD-1 treatment (anti-PD1_Nat.Med, pembrolizumab, 29 patients: 2 ER-/HER2^+^, 15 ER^+^/HER2±, 12 TNBC) was available in the reported literature ^26^. The bulk RNA-sequencing dataset of anti-PD1 treatment in breast cancer was downloaded from NCBI-GEO database with the code: GSE194040 ^27^.

The Cancer Genome Atlas Program-Breast invasive carcinoma (TCGA-BRCA, 1094 tumor tissues and 113 paired normal tissues) dataset was downloaded from GDC Data Portal (https://portal.gdc.cancer.gov/) using the TCGAbiolinks package (2.28.3). The cleaned survival data of the TCGA cohort was obtained from the previous study ^28^. TCGA Pan-Cancer (PANCAN, n=11,057) was downloaded from UCSC Xena (https://xenabrowser.net/datapages/). The expression profiling and clinical data of Molecular Taxonomy of Breast Cancer International Consortium (METABRIC) dataset (2509 primary breast cancer tissues with 548 matched normal tissues) were downloaded from cBioPortal (https://www.cbioportal.org/study/summary?id=brca_metabric) ^29^.

The bulk RNA-sequencing data of Sweden Cancerome Analysis Network-Breast (SCAN-B) dataset (3273 primary breast cancer samples) is available at NCBI-GEO database with the code: GSE96058 (https://www.ncbi.nlm.nih.gov/geo/query/acc.cgi?acc=GSE96058), while the matched clinical data was obtained from the previous literature ^30^. The gene expression profiles of bulk RNA-sequencing with FPKM format were normalized with the log_2_(gene+1) method and duplicated samples were removed. Two publicly available TNBC RNA□seq cohorts were analyzed: the FUSCC□TNBC cohort (n=89) ^31^ and the 2024_Nat.Comm. cohort (n=94) ^32^.

#### 2.1.2 Geneset scoring

The JAK-STAT pathway with 155 genes was retrieved from GSEA database (https://www.gsea-msigdb.org/gsea/msigdb/human/geneset/KEGG_JAK_STAT_SIGNALING_PATHWAY). The core JAKs-STATs pathway genes include JAK1, JAK2, JAK3, STAT1, STAT2, STAT3, STAT4, STAT5A, STAT5B, and STAT6. The immune-stimulating and immune checkpoint molecules, cytotoxic and dysfunctional signatures were reported in previous studies (Table S1) ^33,34^. Above signatures scores were calculated using the AddModuleScore function for single-cell datasets, while single-sample gene set enrichment analysis (ssGSEA) method implemented in GSVA package (version 2.0.2) was employed in bulk RNA-seq datasets.

#### 2.1.3 Clustering and cell annotating

The normal breast epithelial cell dataset was re-annotated using the Seurat package (version 4.4.0) based on following markers (Basal: ACTA2, MYLK, SNAI2; Luminal_progenitor: TNFRSF11A, KIT; Mature_luminal_cell: ESR1, PGR, FOXA1) ^24^.

#### 2.1.4 Pseudotime and cell-cell interaction communication analysis

Monocle (version 2.28.0) and CytoTRACE (version 0.3.3) were applied to predict cell differentiation states and stemness in subclusters (T cells, epithelial cells from tumors and epithelial cells of normal breast) of single-cell datasets. To explore the cellular communications, the samples were divided into Score-High or Score-Low subgroups based on the mean expression of JAK-STAT pathway score in T cells or tumor cells. The cell-cell communication pathway and intensity were inferred using CellChat package (version 1.6.1).

#### 2.1.5 Enrichment analysis

The PROGENy pathway scores of pre-set 13 pathways were calculated using progeny package (version 1.22.0). The differentially expressed genes were identified using the FindAllMarkers function loaded in Seurat (version 4.4.0), with the default threshold: log_2_(fold change) > 0.25 and *p* value < 0.05. The irGSEA package (version 3.3.3) was used for enrichment analysis of differentially expressed genes based on ssGSEA (single-sample gene set enrichment analysis) method. The clusterProfiler (version 4.8.3) package was utilized for GO and KEGG enrichment analysis.

#### 2.1.6 Deconvolution of immune infiltration

For bulk RNA sequencing analysis, immune cell populations were inferred using the CIBERSORT algorithm, which was integrated in the IOBR package (version 0.99.9) with the deconvo_tme function ^35^.

#### 2.1.7 Spatial Transcriptomics

Spatial transcriptomic data were obtained from the GEO database (GSE210616), including 43 10×Visium samples from 14 primary TNBC patients ^36^. Cell type deconvolution was performed using the RCTD package with the scRNA-seq reference dataset GSE176078. Correlation analyses were conducted via the cor.test() function, and spatial gene expression patterns were visualized using Seurat’s SpatialFeaturePlot(). The phenoptr package (version 0.3.2, https://akoyabio.github.io/phenoptr/) was used to analyze cell types spatially adjacent to MIF⁺ tumor cells.

#### 2.1.8 Protein-nucleic docking

Protein sequences were obtained from the UniProt database, and protein-nucleic acid complex structures were predicted using ProteinX, an open-source tool with performance comparable to AlphaFold3. The predicted structures were subjected to energy optimization via the Rosetta Relax module for subsequent computational analysis. The SLC47A1 sequence was retrieved from NCBI (https://www.ncbi.nlm.nih.gov/gene/55244/), and its promoter sequence was defined as the 2000-bp upstream genomic region. The initial 3D model of the protein-promoter complex was predicted using ProteinX, further refined, and exported in PDB format. Flexible docking was performed using RosettaDock, and candidate models were selected based on the built-in Rosetta scoring function. Protein-nucleic acid interactions and binding energies were characterized using LigPlus, FoldX, and PLIP. Conformational representations of the docked complexes were generated using PyMOL.

#### 2.1.9 Predictive Model Construction

A multivariable Cox proportional hazards regression model was constructed to integrate the expression levels of PDCD1, CD274, PDCD1LG2 and the JAK-STAT pathway (JSP) risk score, with the aim of improving prediction of clinical response to immunotherapy in GSE194040 (I-SPY2-990, pembrolizumab, anti-PD-1) and GSE173839 (I-SPY2 trial, durvalumab, anti-PD-L1). A unified model risk score was then calculated for each sample using regression coefficients derived from the coxph() function in the R survival package (version 3.7-0). To evaluate and compare predictive performance, receiver operating characteristic (ROC) curve analysis was performed. The area under the curve (AUC) was used as the primary index to assess the discriminative capacity of individual markers and the combined multi-marker signature. ROC curve visualization and quantitative comparison were conducted using the pROC R package (version 1.18.5).

#### 2.1.10 Survival analysis

The subcluster signatures were defined as the top 50 differentially expressed genes ranked by fold change using the FindAllMarkers function ^37^. Their signature score for each bulk RNA-seq was calculated using the ssGSEA method. The surv_cutpoint function of survminer package (version 0.5.0) was used to determine the optimal cutoff for these these signatures score for subsequent Kaplan-Meier survival analysis with log-rank test in survival package (version 3.7-0).

### 2.2. Experiments

#### 2.2.1 Cell culture

The human triple-negative breast cancer cell line MDA-MB-231 was acquired from Procell System® (CL-0150, Wuhan, China) and confirmed through short tandem repeat (STR) profiling alongside testing for mycoplasma contamination. Cells were cultured in complete RPMI-1640 medium supplemented with 10% fetal bovine serum (10091148, Thermo Fisher Scientific Inc., USA) and 1% penicillin-streptomycin solution (15140122, Thermo Fisher Scientific Inc., USA). MCF10A human normal breast epithelial cells (CL-0525, Procell) were cultured in DMEM/F-12 medium supplemented with 5% horse serum, 20 ng/mL epidermal growth factor (EGF), 0.5 μg/mL hydrocortisone, 10 μg/mL insulin, and 1% penicillin-streptomycin. The full-length STAT4 transcript was cloned into the V5 (EF-1α/GFP & Puro) vector plasmid and verified via Sanger sequencing (Genepharma, Shanghai, China). The plasmid was co-packaged with pGag/Pol, pRev, and pVSV-G systems to produce lentiviral particles in 293T cells. After concentrating the virus and measuring its titer, MDA-MB-231 cells were infected with either the STAT4-overexpressing lentivirus or the control lentivirus for 48 hours. The cells were then subjected to puromycin selection (1 µg/mL) for two weeks to establish a stable cell line expressing target gene.

#### 2.2.2 Cell proliferation assay

CCK-8 cell proliferation assays and colony formation experiments were conducted to assess changes in tumor cell growth phenotypes following gene interference. A total of 3,000 MDA-MB-231 cells from each group were seeded into 96-well plates. At 24-hour intervals, part of the medium was replaced with fresh RPMI-1640 medium containing 10% CCK-8 reagent (HY-K0301, MedChemExpress, China). The cells were incubated at 37°C for 1 hour, and optical density at 450 nm was measured using a spectrophotometer. For the colony formation assay, 1,000 cells were seeded into six-well plates. Once the cells adhered, the culture medium was refreshed every five days. After a 14-day incubation period, the medium was removed, and the cells were fixed with 4% paraformaldehyde. The cells were subsequently stained with 0.1% crystal violet, rinsed with distilled water, and air-dried at room temperature. In the wound-healing assay, scratches were made using a 10 µL pipette tip once cell confluency in the six-well plates reached 100%. After washing the culture medium three times, images were captured. Additional images were taken 12 hours later to observe cell migration, and the migration area was quantified using ImageJ.

#### 2.2.3 Western blotting

Total protein was extracted using RIPA lysis buffer (P0013B, Beyotime, China) and quantified with the BCA Protein Quantification Kit (ZJ102, Epizyme Biotech). A total of 30 µg of protein was subjected to SDS-PAGE for separation, after which the protein bands were transferred to a PVDF membrane. The membrane was blocked with a 5% bovine serum albumin solution at room temperature for 1 hour, followed by incubation with primary antibodies (STAT4: #ET1701-42, dilution ratio = 1:1000, HUABIO; SLC47A1: 20898-1-AP, dilution ratio = 1:2000, Proteintech Group, Inc.; GAPDH: 10494-1-AP, dilution ratio = 1:5000, Proteintech Group, Inc.) overnight at 4°C. Subsequently, a secondary antibody (dilution ratio = 1:5000, SA00001-2, Proteintech) was applied at room temperature for 1 hour. The blots were visualized using an ECL Chemiluminescent Substrate Kit (36222ES60, YEASEN, China) and images were captured with a chemiluminescence imaging system (GelView 6000 Pro, BLT Photon Technology, China).

#### 2.2.4 Real-time qPCR

Total intracellular mRNA was extracted using the TRIzol reagent (15596018CN, Thermo Fisher Scientific Inc.) in accordance with the manufacturer’s instructions. The PrimeScript™ RT reagent Kit with gDNA Eraser (RR047A, Takara) was used to eliminate genomic DNA and reverse-transcribe RNA into cDNA. PCR amplification and fluorescence quantification were conducted using TB Green® Premix Ex Taq™ II FAST qPCR (CN830A, Takara), with fluorescent signals collected on a real-time quantitative PCR instrument (Archimed X4, ROCGENE, China). Relative expression levels were analyzed using the 2^-ΔΔCT^ method. The primer sequences used are as follows: STAT4 (Forward: AGCCTTGCGAAGTTTCAAGA; Reverse: ACACCGCATACACACTTGGA) and GAPDH (Forward: GACTCATGACCACAGTCCATGC; Reverse: AGAGGCAGGGATGATGTTCTG).

#### 2.2.5 RNA-sequencing and analysis

Total RNA was extracted using TRIzol reagent, and its concentration and integrity were assessed with a NanoDrop 2000 spectrophotometer and an Agilent 2100/LabChip GX instrument. Following library construction, sequencing, and quality control of FASTQ files, HISAT2 and StringTie software were utilized to align the data to the reference genome and construct transcripts. Differential gene expression was determined from the expression matrix with fragments per kilobase of transcript per million fragments mapped (FPKM) format, employing the Limma package (version 3.56.1) in R with thresholds set at p < 0.05 and log2(fold change) > 1. Gene enrichment analysis was conducted using the clusterProfiler (version 4.8.2) and GSVA (version 1.48.3) packages, while visualization was carried out using the ggplot2 (version 3.5.1) and pheatmap (version 1.0.12) packages.

#### 2.2.6 Animal experiment

The animal experimental protocol was approved and overseen by the Medical Ethics Committee of Sichuan University (ID: K2024019). Female nude mice, procured from GemPharmatech Co., Ltd., were maintained in a specific pathogen-free (SPF) animal facility at the West China School of Pharmacy, Sichuan University, with unrestricted access to food and water. A total of 2 × 10^6 MDA-MB-231 cells were injected subcutaneously into the dorsal region of each mouse, with a minimum of five mice per group. Tumor volume was assessed weekly, and after three weeks, the mice were euthanized. Tumor tissues were excised, measured, and prepared for subsequent histological staining analysis.

#### 2.2.7 Immunohistochemistry (IHC)

Fresh tumor tissues were fixed in 10% neutral formaldehyde for 24 hours, followed by gradient dehydration, and then embedded in paraffin to create paraffin blocks. Tissue sections, 5 µm in thickness, were cut and subsequently deparaffinized and rehydrated. Peroxidase activity was inhibited using 3% hydrogen peroxide, followed by blocking with 5% bovine serum albumin (BSA). The primary antibodies (STAT4: 13028-1-AP, dilution ratio = 1:200, Proteintech Group, Inc.; SLC47A1: bs-9284R, dilution ratio = 1:500, Bioss Inc.) were then applied and allowed to incubate overnight at 4°C. The sections were subsequently incubated at room temperature with horseradish peroxidase (HRP)-conjugated anti-rabbit IgG (SV0004, Boster) and developed using a DAB kit (AR1027-1, Boster). Image analysis was conducted using Image-Pro Plus 6.0 (Media Cybernetics, Rockville, USA), with integrated optical density (IOD) values calculated accordingly.

### 2.3. Statistics analysis

All bioinformatics analyses were conducted in R software (4.3.0), while the statistical analysis of experimental results was performed using GraphPad Prism 9, presented as mean ± standard deviation (SD). Bioinformatic results were visualized in R software (4.4.0) using ggplot2 (3.5.1) for boxplot, pheatmap (1.0.12) and ComplexHeatmap (2.22.0) for heatmap. Linear correlation analysis was computed using the Spearman’s algorithm. The statistics significance of non-normally distributed data was calculated using Wilcoxon Signed Rank Test for two subgroups and Kruskal-Wallis for multiple-subgroups comparison. For normally distributed data, the unpaired T-test was used to compare the two groups, while significant differences among multiple groups were assessed using a one-way ANOVA method. A p-value of less than 0.05 was considered statistically significant.

## 3. Results

### 3.1 Dynamic swift of JAK-STAT pathway in carcinomas

In the TCGA pan-cancer cohort, JAK-STAT pathway (JSP) is typically higher in adjacent non-tumor tissues than in tumor tissues, and patients with elevated JSP expression generally experience poorer prognoses. Interestingly, in breast cancer, patients with high JSP scores display better survival outcomes, which challenges the conventional view of JSP as an oncogenic pathway (Fig S1). This indicates a heterogeneous role for the JSP pathway in breast cancer. We next analyzed the primary breast cancer single-cell dataset (GSE176078, 26 cases) to examine the expression patterns and dynamic features of the JSP at single-cell resolution. JSP scores were calculated using the ssGSEA method, which revealed predominant expression in T cells (Fig. 1B-E). Further annotation indicated elevated JSP scores within NK cells, CD4^+^ T cells, and CD8^+^ T cells (Fig. 1F). Using the Ro/e algorithm, we assessed the preferential distribution of JSP across various subpopulations, finding that cells with high JSP scores were primarily enriched in T cells, while paracancerous epithelial and tumor cells showed significantly lower enrichment (Fig. 1G-H). Pseudotime analysis illuminated the developmental trajectory from paracancerous epithelial cells to tumor cells (Fig. 1I-L, Fig.S2B). During this progression, JSP demonstrated a dynamic shift, initially decreasing and then increasing, suggesting differential biological functions and dynamic dependencies between paracancerous and tumor epithelial cells (Fig. 1L), which is consist with previous reports: basal cells differentiated into luminal cells in breast cancer (STAT3 serves as a crucial transcription factor in basal cells) ^38^; tumorigenesis from basal cells to luminal cells depended STAT3 in prostate cancers ^39^. Progeny pathway activity analysis corroborated this trend, underscoring greater JSP activity in epithelial cells and heterogeneity during tumor cell differentiation (Fig. 1M). These findings demonstrate that the JAK-STAT pathway exhibits heterogeneous expression patterns and dynamic transitions within the breast tumor cells and epithelial cells.

### 3.2 JAK−STAT pathway facilitates tumor malignant phenotypes and immunosuppression

Next, we analyzed the potential biological functions of the JAK-STAT pathway in tumor cells and its interplay with the immune microenvironment. Within tumor cell subgroups, JSP was enriched in the LumA and LumB subtypes (Fig. 2A). Correlation analysis revealed a positive association between JSP and various cancer characteristic scores, including proliferation, invasion, and differentiation (Fig. 2B). By classifying tumor cells into JSP-high and JSP-low groups based on median JSP scores, it was observed that JSP-high tumor cells exhibited a higher degree of malignancy, along with elevated expression of these characteristic scores (Fig. 2C). Subsequently, we performed differential gene analysis to uncover potential biological functions of JSP in tumor cells. Oncogenes, such as MYC, IL6ST, and NEAT1, were found to be upregulated in JSP-high tumor cells (Fig. 2D). Pathway enrichment analysis indicated that these differential genes are enriched in oncogenic signaling pathways like MAPK, NF-κB, and PD-L1/PD-1 immune checkpoints (Fig. 2E). Cell communication analysis revealed differences in immune microenvironment signaling intensity between JSP-high and JSP-low patients (Fig. 2F). Signals like IL6, CCL, EGF, and IL10 were present only in JSP-high patients (Fig. 2G). Moreover, enhanced immunosuppressive signaling such as MIF-CD74+CXCR4 and MIF-CD74+CD44 was observed in JSP-high patients (Fig. 2H). In addition, we verified the spatial co-localization of MIF-expressing tumor cells and T cells using the breast cancer spatial transcriptomic dataset GSE210616 (Fig S3). These findings expand our understanding of the oncogenic role of JSP in breast cancer and its interplay within the immune microenvironment. JSP resonates with carcinogenic signals like MAPK and NF-κB and is involved in immunosuppression through MIF-CD74+CXCR4 and MIF-CD74+CD44 signaling pathways.

**Figure 2.**
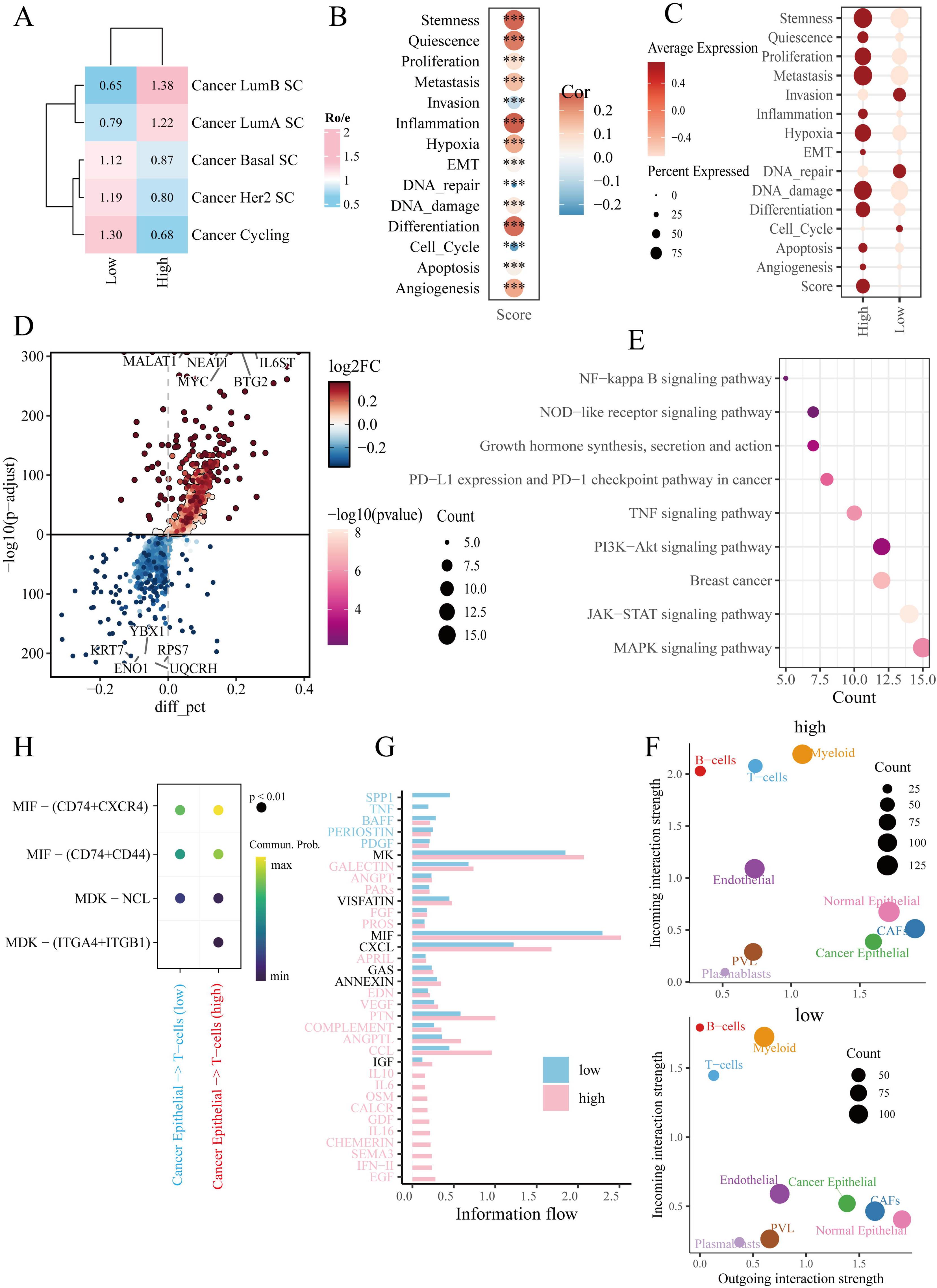
The JAK-STAT pathway promotes tumor malignant phenotypes and immunosuppression. (A) Ro/e index analysis demonstrates the preferences of JSP-high versus JSP-low cells in tumor cells. (B) Correlation of JSP scores with 14 cancer-related phenotype features. (C) Expression of cancer phenotype signatures in JSP-high and JSP-low tumor cells. (D) Differentially expressed genes in JSP-high versus JSP-low tumor cells. (E) KEGG enrichment analysis of differentially expressed genes. (F) CellChat analysis illustrating the strength of cellular communication in JSP-high and JSP-low samples. (G) Bar chart of CellChat signaling flows. (H) Cell-to-cell communication between cancerous epithelial cells and T cells.

### 3.3 JAK-STAT pathway is essential for breast epithelial cell differentiation

Given the differential expression patterns of JSP in adjacent non-cancerous and tumor epithelial cells (Fig. 1D-F), we analyzed another human normal breast epithelial single-cell dataset (GSE161529) to explore the role of JSP in breast epithelial cell differentiation under normal physiological conditions. JSP showed higher expression in basal epithelial cells (Fig. 3A-C). Pseudotime analysis revealed the differentiation trajectory of epithelial cells: from luminal progenitor to mature luminal and basal cells (Fig. 3D, Fig S4A). During epithelial cell differentiation, JSP expression increased (Fig. 3E). In JSP-high epithelial cells, enhanced activity of the p53 and TGFβ pathways was observed (Fig. 3F). Differential gene analysis and GO enrichment for JSP-high versus JSP-low epithelial cells indicated that JSP signaling is involved in maintaining normal breast epithelial cell functions, such as focal adhesion, vesicle lumen, and secretory granule lumen (Fig. 3G-H). KEGG pathway analysis suggested that JSP is associated with several important signaling pathways in epithelial cells, including Estrogen, TGFβ, and FoxO pathways (Fig. 3I). These results suggest that, under physiological conditions, the normal development and functional maintenance of breast epithelial cells depend on JSP signaling.

**Figure 3.**
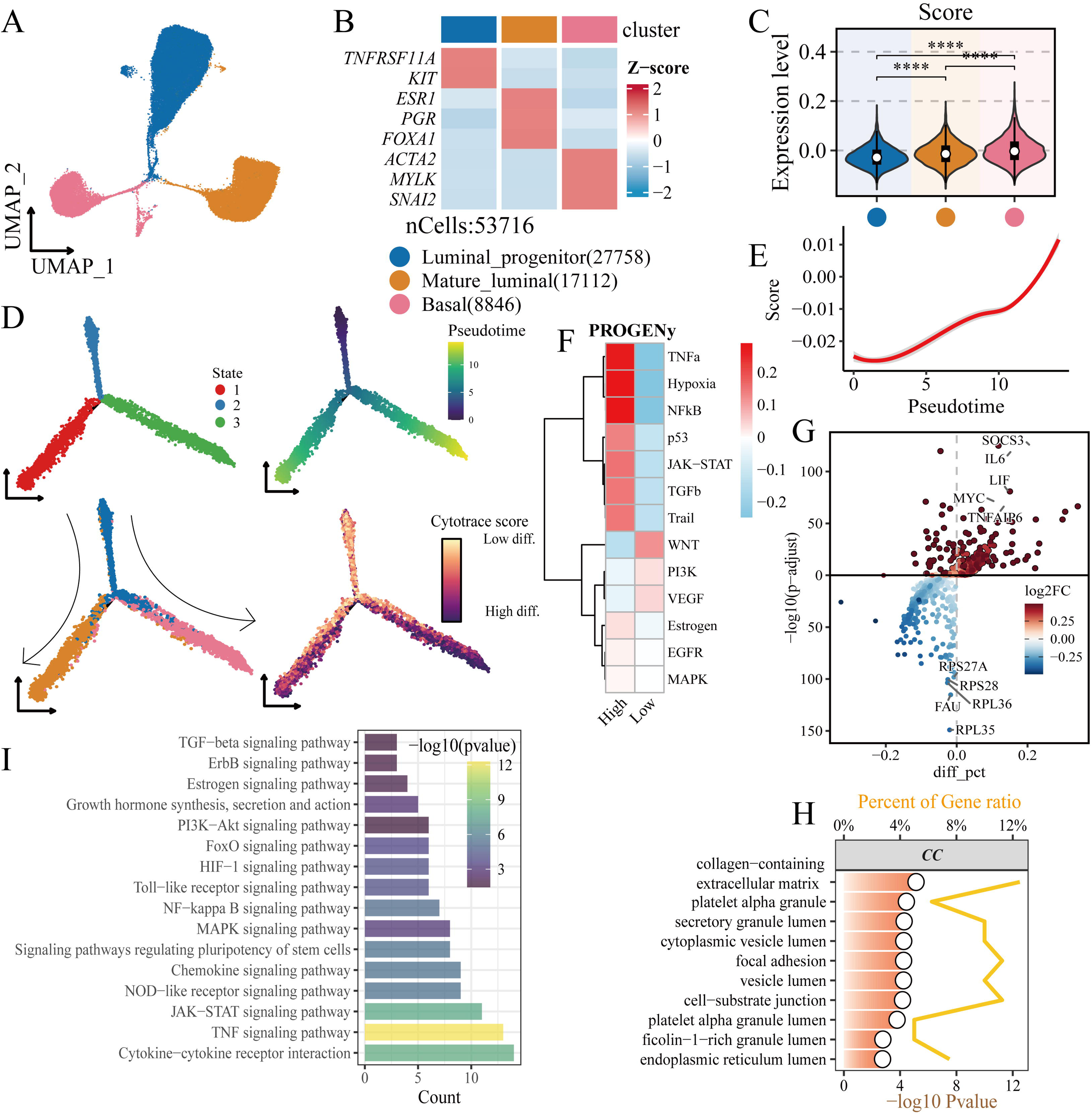
JAK−STAT pathway is essential for normal breast epithelial cell differentiation. (A) UMAP plot of annotated normal epithelial cells. (B) Core markers of mammary epithelial cells. (C) Expression levels of JSP score across mammary epithelial cell subgroups. (D) Pseudotime trajectory analysis using Monocle2. (E) Relationship between JSP expression and pseudotime scores. (F) Progeny pathway activity, with epithelial cells categorized into high and low groups based on median JSP expression. (G) Volcano plot of differentially expressed genes. (H-I) GO and KEGG enrichment analyses of differentially expressed genes. Significance: *, *p* < 0.05; ** *p* < 0.01; ***, *p* < 0.001.

### 3.4 JAK−STAT pathway mirrors anti-tumor immunity activity of T cells

Given the highest expression of JSP in T cells (Fig. 1E, F), we explored the potential biological functions of JSP in T cells. Major and minor annotations showed elevated JSP scores in CD4^+^ T, CD8^+^ T, and NK cells (Fig. 4A-B). T cells were divided into JSP-high and JSP-low subgroups based on median JSP scores, with no significant differences in T cell composition between the two groups (Fig. 4C). Since the number of T cells remained unchanged, we hypothesized that JSP might impact T cell state and function. We collected representative T-cell state signatures to comprehensively assess the status and activity of major T cell subgroups (Fig. 4D) ^40^. JSP-high T cells exhibited increased functional scores (such as TCR signaling, IFN-γ response, cytotoxicity) and decreased metabolic scores (glycolysis, fatty acid metabolism, oxidative phosphorylation) and exhaustion scores (Fig. 4E-F). Further analysis of representative functional genes in CD8^+^ T cells confirmed this, showing higher expression of T cell cytotoxic factors (GZMB, TBX21, IL2RA) and lower expression of T cell exhaustion markers (PDCD1, KLRC1, CTLA4, LAG3, TIGIT) in the JSP-high group (Fig. 4G-H). Additionally, pseudotime analysis indicated that JSP plays an important role in promoting T cell differentiation (Fig. 4I-J, Fig S4B). Differential gene analysis revealed that JSP-high T cells exhibit higher expression of T cell antitumor factors (IFNG, TNF: encoding TNF-α) and immune activation-related receptors (IL2RA: CD25, IL10RA) (Fig. 4K). Pathway enrichment suggested that these differential genes are enriched in T cell activation and differentiation pathways (Fig. 4L). These findings collectively indicated that T cells with high expression of the JAK-STAT pathway have stronger antitumor activity.

**Figure 4.**
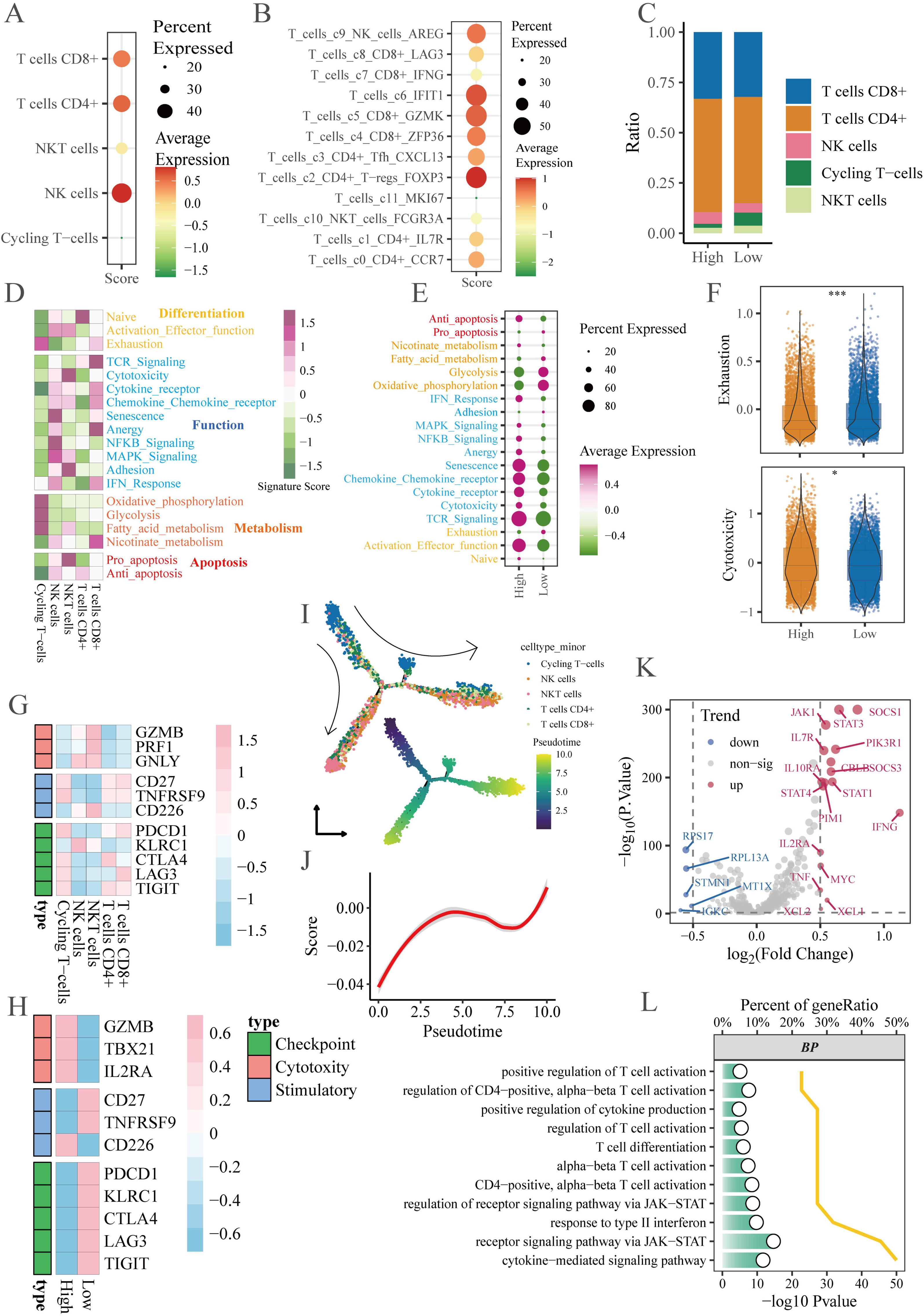
JAK−STAT pathway is associated with enhanced T cell function. (A) JSP scores across major T cell subgroups in GSE176078. (B) JSP scores across minor T cell subgroups. (C) Proportions of T cell subgroups. T cells were classified into high and low groups based on the average JSP score. (D) T cell state scores of major T cell subgroups. (E) T cell state scores based on JSP grouping. (F) Related to (E), exhaustion and cytotoxicity scores in T cells. (G-H) Heatmaps of representative functional genes in major T cell subgroups and JSP subgroups. (I) Monocle2 pseudotime analysis UMAP plots; the top panel shows cell types, the bottom panel displays pseudotime scores. (J) Line plot showing the relationship between JSP score and pseudotime score. (K) Differential genes between T cell JSP subgroups. (L) GO enrichment analysis of differential genes between JSP-high and JSP-low T cells. Significance: *, *p* < 0.05; ** *p* < 0.01; ***, *p* < 0.001.

### 3.5 JAK−STAT pathway enhances immunotherapy sensitivity and predicts immunotherapy response

Given that JSP is associated with T cell cytotoxic activity, we investigated the relationship between JSP and immunotherapy response, as well as its clinical significance. In the bulk RNA-seq immunotherapy cohort (GSE194040), JSP levels were higher in responders, accompanied by an increase in immune markers and a higher proportion of TNBC samples (Fig. 5A-B). JSP scores also demonstrated a relatively accurate prediction of immunotherapy response with ROC=0.7 (Fig. 5C-D). To further expand the clinical applicability of the JSP score, we combined it with the mRNA expression levels of PD-L1/PD-1, which significantly improved the predictive accuracy for immunotherapy response, with the AUC exceeding 0.8 (Fig S5A). Subsequently, we further explored the significance of JSP before and after immunotherapy in a single-cell immunotherapy dataset (anti-PD1_Nat.Med) (Fig. 5E-K). JSP was primarily expressed in T cells (Fig. 5E-F) and enriched in TNBC samples (Fig. 5G). In JSP-high patients, there was a higher proportion of TNBC cases and T cell expansion, along with improved immune infiltration, characterized by increased proportions of T, B, and pDC cells, and a decrease in cancer cells and fibroblasts (Fig. 5H). JSP showed a positive correlation with T, B, and pDC cell proportions, and markedly increased after immunotherapy (Fig. 5I-J). Additionally, JSP was able to predict T cell expansion more accurately compared to other immune cell proportions (Fig. 5K). These results demonstrated that patients with high JAK-STAT pathway level exhibit a greater response rate to immunotherapy, especially in TNBC.

**Figure 5.**
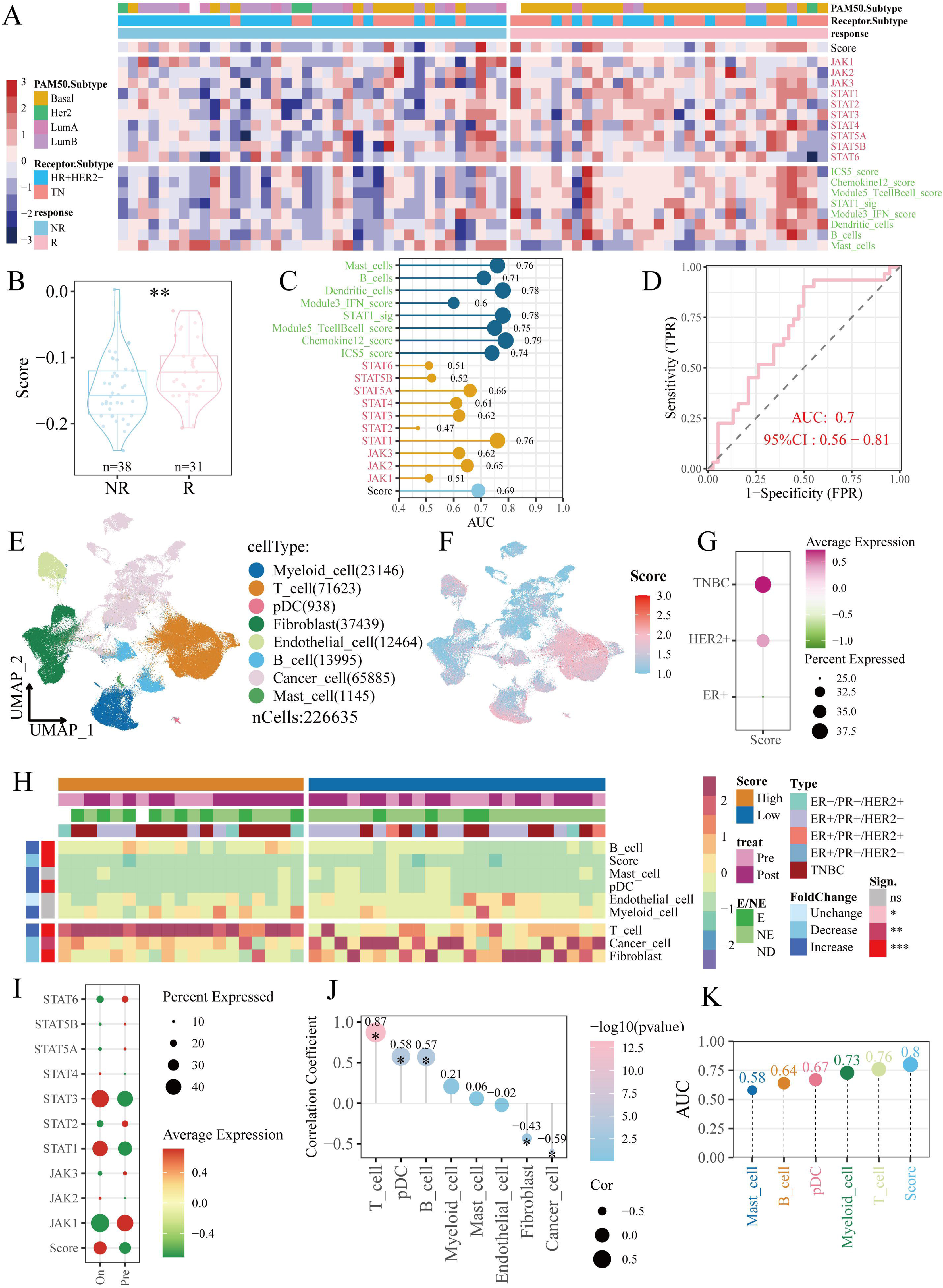
JAK−STAT pathway enhances immunotherapy sensitivity and predicts immunotherapy response. (A) Expression patterns of JSP in the bulk RNA-seq immunotherapy cohort: GSE194040. Red text indicates 10 core JSP components, while green text shows 8 immune markers. TN, triple-negative breast cancer; Score, JSP score. (B) JSP scores in immunotherapy responders (R) versus non-responders (NR). (C) Area Under Curve (AUC) of Receiver Operating Characteristic (ROC) curve for predicting immunotherapy response. (D) ROC curve for JSP predicting immunotherapy response extracted from (C). (E) UMAP of the single-cell dataset: breast cancer anti-PD1 immunotherapy cohort (anti-PD1_Nat.Med). (F) JSP score in UMAP, pDC, plasmacytoid dendritic cells. (G) Distribution of JSP scores across clinical subtypes of anti-PD1_Nat.Med cohort. (H) Immune microenvironment characteristics (scaled cell type ratio) of JSP-high versus JSP-low patients. (I) Expression of JSP and core JSP components before and during immunotherapy (Pre vs. On). (J) Correlation between JSP scores and cell proportions. (K) ROC curve AUC values: Prediction of T cell expansion using JSP scores and cell composition proportions. Significance: *, *p* < 0.05; ** *p* < 0.01; ***, *p* < 0.001.

### 3.6 STAT4 promotes the proliferation of cancer cells and epithelial cells

Given the heterogeneous expression pattern in tumor cells, epithelial cells and T cells, we obtained these cell-type specific genes from pseudotime trajectory analysis (Fig. 6A, Fig S2B, Fig.S4). Tumoral JSP score was negatively associated with patient survival, while epithelial and T cell JSP score were related to favorable prognosis (Fig. 6A). Given STAT4, one of the core JAK-STAT pathway numbers, performed similar expression pattern with JAK-STAT pathway (high expression is associated with favorable prognosis and is primarily observed in T cells) (Fig S5B-C, Fig S6), we selected STAT4 as an interest gene to explore its role in tumor cells and epithelial cells. In MDA-MB-231 breast cancer cells, overexpression of STAT4 promoted proliferation, tumor growth, migration and invasion (Fig. 6C-E, Fig.S7A-B). Similar results also were observed in normal epithelial MCF10A cells (Fig. 6F-H). RNA-sequencing of MDA-MB-231 cells indicated that STAT4 enhances G2M checkpoint and mitotic pathways (Fig. 6L). STAT4 co-activates the JAK-STAT and WNT pathways (Fig. 6M). KEGG enrichment analysis suggests that STAT4 is involved in the ferroptosis pathway. Given that STAT4 is a potential upstream transcription factor of the ferroptosis inhibitor SLC47A1 ^41,42^, we confirmed that STAT4 upregulates SLC47A1 in breast cancer both in vivo and in vitro (Fig S7). STAT4 directly binds to the SLC47A1 promoter to promote its transcription (Fig S8). Altogether, these results implied that STAT4, as a key transcriptional orchestrator of JAK-STAT pathway, facilitated malignant and epithelial cell proliferation and function.

**Figure 6.**
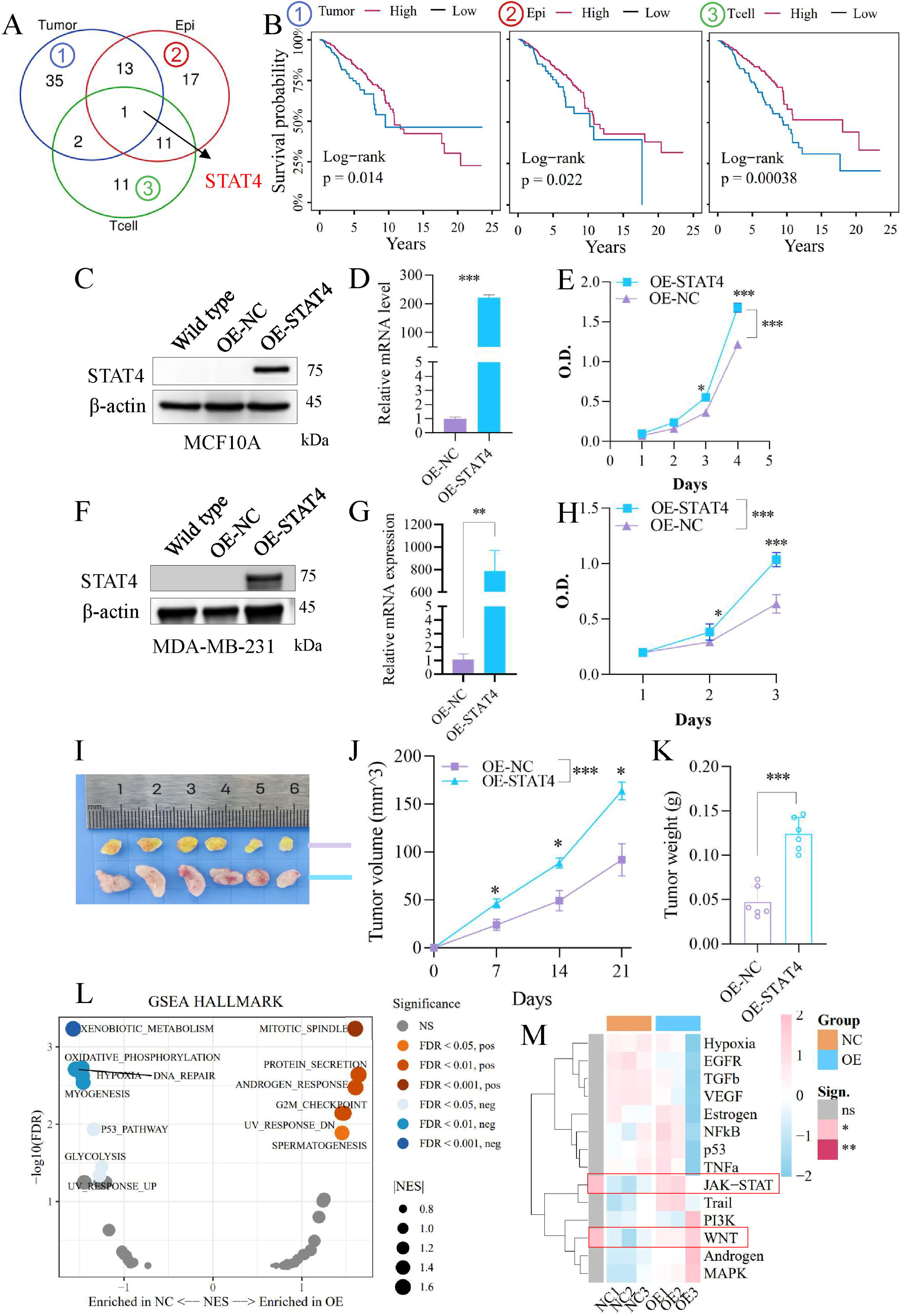
STAT4 promotes tumor cell and normal epithelial proliferation. (A) Venn diagram displaying the JAK-STAT pathway signature genes specific to tumor cells, epithelial cells, and T cells, which were derived from pseudotime trajectory analysis using Monocle2. (B) Survival analysis of the above cell-type-specific JAK-STAT pathway genes in the TCGA dataset. (C-E) Overexpressing STAT4 in normal breast epithelial MCF10A cells, detected by Western blotting (C) and qPCR (D). (E) MCF10A cells were assigned to OE-NC (control) or OE-STAT4 groups for cell proliferation assay. (F-H) Overexpressing STAT4 in MDA-MB-231 breast cancer cells. (F) Protein levels of STAT4 following overexpression. (G) STAT4 mRNA levels following overexpression. (H) Cell proliferation assessed by CCK-8 assay. (I-K) Xenograft tumor experiments in nude mice: tumor volume (J) and weight (K). (L) Gene Set Enrichment Analysis (GSEA) of RNA-sequencing after STAT4 overexpression. (M) Progeny pathway activity analysis. Significance: *, *p* < 0.05; ** *p* < 0.01; ***, *p* < 0.001.

## 4. Discussion

We analyzed single-cell datasets to explore the intra-tumoral heterogeneity of the JAK-STAT pathway. Its expression was highest in T cells and lowest in epithelial cells, with normal epithelial cells expressing higher levels than tumor cells. In all epithelial cells containing normal epithelial and tumor cells, JAK-STAT signaling was reprogrammed during basal-to-luminal differentiation and tumor formation, showing an initial decrease followed by an increase. Its activity declines during transition from normal epithelium to early tumors, then rises in advanced tumor stages due to tumor subtype heterogeneity and subtype-specific dependency. In tumor cells, higher JAK-STAT activity correlated with enhanced pro-tumor functions (e.g., migration, invasion, proliferation) and immunosuppressive signaling through the MIF-CD74 axis. Recent studies identify a conserved MIF□CD74/CD74⁺ lipid-associated macrophages (LA-MAMs) axis driving T□cell exhaustion and multi□organ metastasis in breast cancer, correlating with poor survival ^43^. Our findings reveal that tumor□intrinsic JAK□STAT signaling may depend on similar immunosuppressive cascade, while STAT4 activation in T□cell reverses immune suppression. Combining MIF□CD74 blockade with targeted STAT4 activation may synergistically remodel the metastatic niche and improve immunotherapy efficacy in TNBC. To exclude tumor background, we used single-cell data from normal human mammary epithelial cells to assess the role of JAK-STAT signaling in differentiation, finding that epithelial differentiation depends on JSP, potentially working with TNF and TGFβ signaling to maintain luminal integrity and secretory function. Meanwhile, JSP is involved in T cell differentiation, representing enhanced cytotoxicity and functionality of terminal and effector-like T cells. To apply our findings, in a bulk RNA-seq immunotherapy cohort, higher JSP levels correlated with better treatment response, increased immune markers, with accessible accuracy for response prediction. Single-cell analysis confirmed JSP enrichment in T cells, with enhanced immune infiltration and T cell expansion after post-immunotherapy. In brief, High JSP levels predict better immunotherapy response, particularly in TNBC. Our findings suggest that baseline JAK-STAT pathway activity could serve as a potential biomarker for predicting the efficacy of PD-1 immunotherapy in breast cancer. This discovery holds promise for advancing precision medicine by improving response rates and reducing treatment-associated toxicity ^44^. In survival profiling, higher T-cells- and normal epithelial-specific JSP scores correlate with favorable patient survival, whereas elevated tumor-intrinsic JSP scores are associated with poor prognosis. This can be attributed to the predominant expression of JSP in T cells, which enhances T cell-mediated anti-tumor immunity and counterbalances its pro-tumor effects within cancer cells.

STAT4 exhibits contrasting roles in tumor cells and immune cells, highlighting its dual nature in cancer biology. On one hand, STAT4 in tumor cells drives carcinoma progression. For example, STAT4 has been shown to transactivate KDM5D, which erases H3K4me2/3 and H3K27ac marks, leading to the repression of cell-cell junction genes and antigen presentation genes. This, in turn, facilitates cancer metastasis and immune evasion in colorectal cancer ^45^. Additionally, STAT4 contributes to the progression of triple-negative breast cancer by co-transactivating with STAT3 to establish the IL12R/JAK2-STAT3-STAT4/PD-L1 axis ^23^. We also confirmed STAT4 facilitated cancer progression through orchestrating with JAK-STAT pathway and involved ferroptotic resistance through upregulating SLC47A1. On the other hand, STAT4 plays a critical role in maintaining the anti-tumor efficacy of immune cells within TME. Recognized as a favorable prognostic biomarker in breast cancer ^23,46,47^ and gastric cancer ^48^, STAT4 promoted anti-tumor immunity mediated by Th1, Th17 through increasing IFN-γ production in HNSCC ^49^. It also enhanced NK and NKT cell cytotoxicity, promoted Th1 cell proliferation and differentiation with a IL12-dependent manner ^50–52^, and strengthened IFN-γ-mediated macrophage-T cell crosstalk ^49,53^. Here, our bioinformatics analysis and experimental findings further reinforce the aforementioned perspective: STAT4 accelerates tumor cell progression and normal epithelial proliferation, while in T cells, it serves as an indicator of enhanced T-cell cytotoxicity.

Preclinical studies identified the potential of targeting JAK-STAT pathway for cancer treatment through small molecular inhibitors ^54–57^. Furthermore, a series of clinical trials unveil that JAK or STAT inhibitors perform synergetic potential towards anti-PD1 immunotherapy in cancers ^54,58^. For instance, STAT inhibitors (BBI608, Fludarabine, Celecoxib) or JAK inhibitors (Ruxolitinib, Itacitinib) combined with PD-1/PD-L1 inhibitors (Pembrolizumab, Nivolumab, Toripalimab) for phase 1/2 clinical trials ^58^. Meanwhile, STAT3 hinders CD8^+^ T cell recruitment to tumor microenvironment through suppressing IFNγ-CXCL10 (chemoattractant for T cells)-CXCR3 (CXCL10 receptor) signaling in lung cancer preclinical models, which implied targeting STAT3 in T-cell also could enhance T cell infiltration, even immunotherapy efficiency ^59,60^. Moreover, STAT3 is one of the upstream transcriptional factors of CD274 (coding PD-L1) to involve PD-L1-mediated immune evasion ^61,62^. Targeting JAK/STAT-PD-L1 crosstalk is regarded as a druggable strategy to boost immunotherapy in lung cancers ^57^. JAK1 inhibitor (ruxolitinib) improved anti-PD-1 treatment response through reshaping T cell differentiation in a lung cancer cohort ^63^, while ruxolitinib reprogramed exhausted T cells in Hodgkin’s lymphoma ^64^. However, the strategy of targeting the JAK-STAT signaling pathway for breast cancer treatment has proven to be both underwhelming and disappointing, with the number of related clinical trials lagging behind those for other types of cancer ^65–67^. JAK1/2 inhibitor Ruxolitinib in metastatic TNBC cohort didn’t reach efficacy endpoint without combination with immunotherapy ^68^. In a clinical trial (NCT01594216) evaluating the combination of Ruxolitinib and exemestane, a steroidal aromatase inhibitor, for the treatment of HR^+^ metastatic breast cancer, none of the patients achieved complete or partial response. Additionally, nearly 50% of the participants experienced anemia as an adverse effect ^69^. These findings suggest that Ruxolitinib has limited utility for treating advanced breast cancer. Another clinical trial (NCT06731153) is currently underway to investigate whether the JAK1 inhibitor Ivarmacitinib can overcome resistance to PD-1 inhibitor (Camrelizumab) immunotherapy in TNBC. In a clinical trial (NCT03195699) evaluating the efficacy of the oral STAT3 inhibitor tinengotinib (TTI-101) as a monotherapy for advanced metastatic solid tumors, including 7 cases of breast cancer, 12% of patients achieved a partial response. Among HCC patients, the overall response rate (ORR) was 18%. However, adverse events or side effects of varying degrees were observed in 98% of the participants ^70^. Meanwhile, considering the critical role of the JAK-STAT pathway in immune cells, its inhibition can lead to immunosuppression and immune cytotoxicity ^17,71^. For instance, JAK inhibitors (such as ruxolitinib) impair the anti-tumor functions of DCs ^72^, NK cells ^73^, and T cells ^74^. Our analysis also partially revealed why the effectiveness of JAK/STAT inhibitors in breast cancer treatment remains unsatisfactory: broad inhibition of the JAK-STAT pathway may hinder T-cell activation and compromise T-cell-mediated anti-tumor cytotoxicity; JAK-STAT pathway is also crucial for normal epithelial differentiation. Considering the complexity and context-specific nature of JAK-STAT pathway signaling and its transduction network within TME ^17^, leveraging the JAK-STAT pathway as a biomarker for immune therapy response is a practical and effective approach. Based on the above clinical evidence and our findings, administering JAK-STAT inhibitors before or concurrently with immunotherapy may impair T-cell cytotoxicity and disrupt normal epithelial differentiation in breast cancer patients. Instead, sequential delivery of JAK inhibition following immunotherapy represents a promising immune-sensitizing strategy, particularly for the TNBC subtype.

Recent studies have highlighted the critical role of selecting optimal therapeutic windows for JAK inhibitors (JAKi) in immunotherapy. In patients with non-small cell lung cancer (NSCLC), intermittent treatment of JAK inhibitors (JAKi) during anti-PD-1 immunotherapy, rather than continuous combination therapy, has demonstrated potential for improving treatment outcomes. This approach reshapes T cell differentiation, characterized by an increase in memory and effector-memory CD8^+^ T cells and a reduction in exhausted CD8^+^ T cells ^63^. Another study in Hodgkin lymphoma found that adding JAK inhibitors at the late stage of anti-PD-1 therapy similarly increased the proportion of cytotoxic CD8^+^ T cells, accompanied by a reduction in pro-tumor neutrophils and MDSCs ^64^. These findings suggest that the therapeutic window should also be carefully considered when introducing JAK inhibitors into breast cancer treatment, even immunotherapy. A Phase II clinical trial (NCT06731153, not yet recruiting) aims to evaluate whether the JAK1 inhibitor Ivarmacitinib combined with the PD-1 inhibitor Camrelizumab can overcome immunotherapy resistance in triple-negative breast cancer. Other clinical trial (NCT05384119) targeting palbociclib-resistant HR+/HER2- breast cancer by combining the STAT3 inhibitor TTI-101 with established therapies to assess its safety, tolerability, and efficacy has concluded. Additionally, immune-activating cytokine IL-12 binds to its receptors IL12R-β1/β2, triggering Tyk2/Jak2 signaling to drive STAT4 phosphorylation, dimerization and nuclear translocation; this further upregulates IFN-γ transcription to potentiate T-cell and NK-cell-mediated antitumor immune responses ^75,76^. Several preclinical evidences have identified the efficacy of IL12 against TNBC ^77,78^. These findings suggest that combining IL□12-mediated STAT4 activation with STAT3 inhibition may open a new avenue for immunotherapy in breast cancer. Some unresolved questions include: Does JAK-STAT inhibition have therapeutic value in breast cancer immunotherapy? How can a reasonable therapeutic window be designed? Which breast cancer subtypes should be selected for further clinical trials? Is there a difference between JAK inhibition and STAT inhibition? Which is more effective? Further experiments are needed to address these questions.

In our analysis of the immunotherapy cohort, only PD-1 inhibitor datasets were included. This may limit the representativeness of our findings with respect to other ICIs, such as PD-L1 inhibitors and CTLA-4 inhibitors. Additionally, given one of the obvious molecular characteristics of TNBC subtype: MES (elevated JAK/STAT3 singling pathway ^31,79^) is the abnormal activation of JAK-STAT signaling, our profiling not refine and investigate this pathway based on these subtypes because lack of subtype-paired single-cell cohorts. Notably, the IM subtype displayed elevated JAK-STAT pathway expression and activity (Fig S5B-C), highlighting meaningful therapeutic implications for patients harboring this subtype. In addition, the JAK-STAT pathway gene set may need refinement to better reflect cancer type dependency. In summary, our results emphasize the JAK-STAT pathway’s heterogeneity in epithelial, tumor, and T cells, with patients exhibiting high pathway activity gaining more substantial benefits from immunotherapy.

## 6. Author Contributions

J.Z.: conceptualization, bioinformatics analyses, experimental work, and writing of the original draft. H.Z.: experimental support and data acquisition. L.Y. and F.P.: supervision, funding acquisition, project administration, and manuscript review and revision.

## 7. Acknowledgements

We thank all the editors and reviewers for their efforts and insightful comments.

## 9. Data availability

All clinical datasets are publicly available and stated in the methods section. All R codes are available on GitHub (https://github.com/Jianbo1999/2025-JAK-STAT-breast-cancer).

## 10. Ethnics statement

Animal experiments were approved by the Medical Ethics Committee of Sichuan University (ID: K2024019).

---

**Figure S1.**
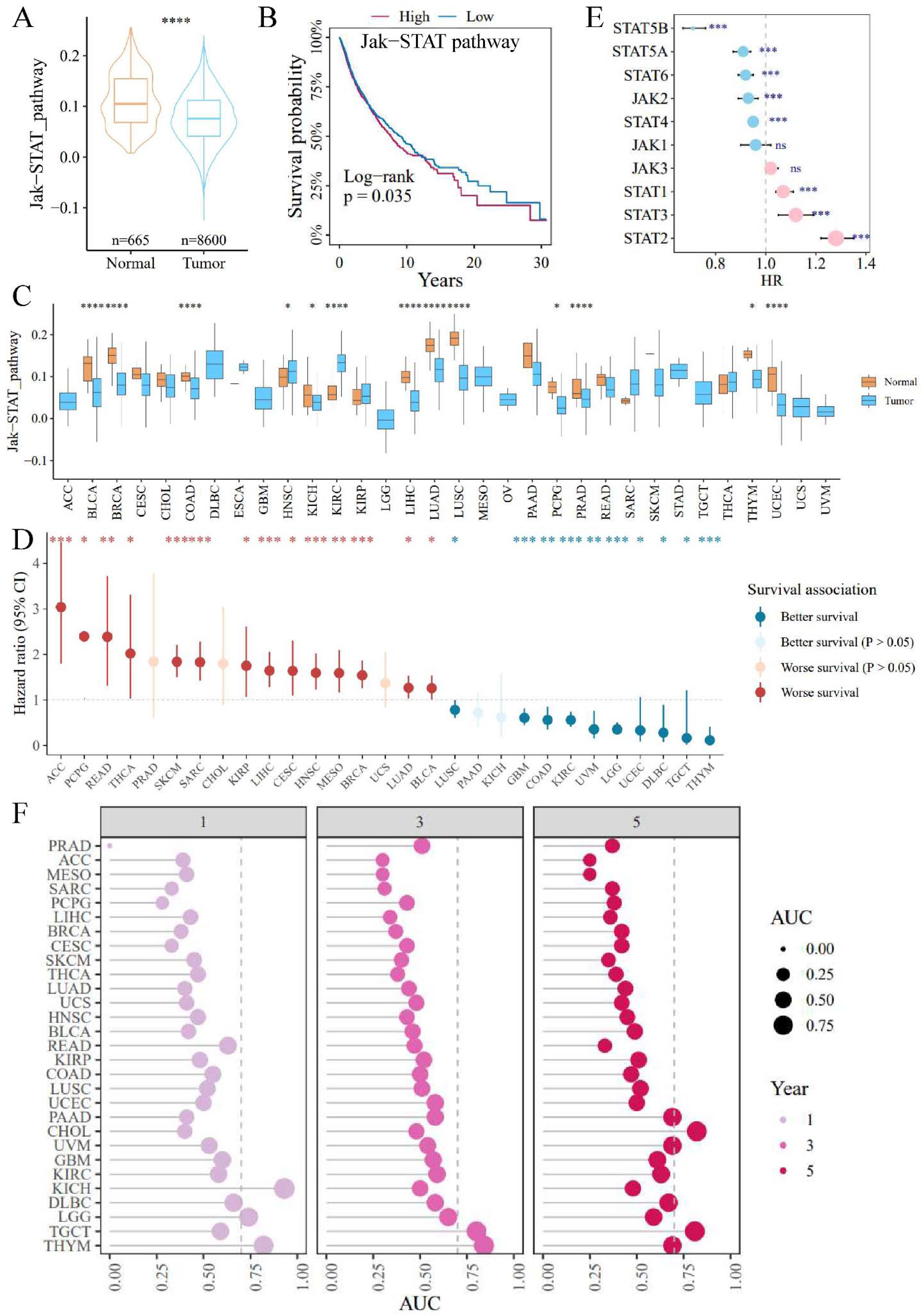
The JAK-STAT signaling pathway is abnormally expressed and prognosis-related in cancers. (A) Expression of JSP in adjacent non-tumor and tumor tissues across a pan-cancer cohort. (B) Survival curves based on JSP pathway expression, stratified by median value. (C) Expression of JSP in adjacent non-tumor and tumor tissues within specific cancer types. (D) Univariate COX analysis across different cancer types. (E) Univariate COX analysis of core JSP components.

**Figure S2.**
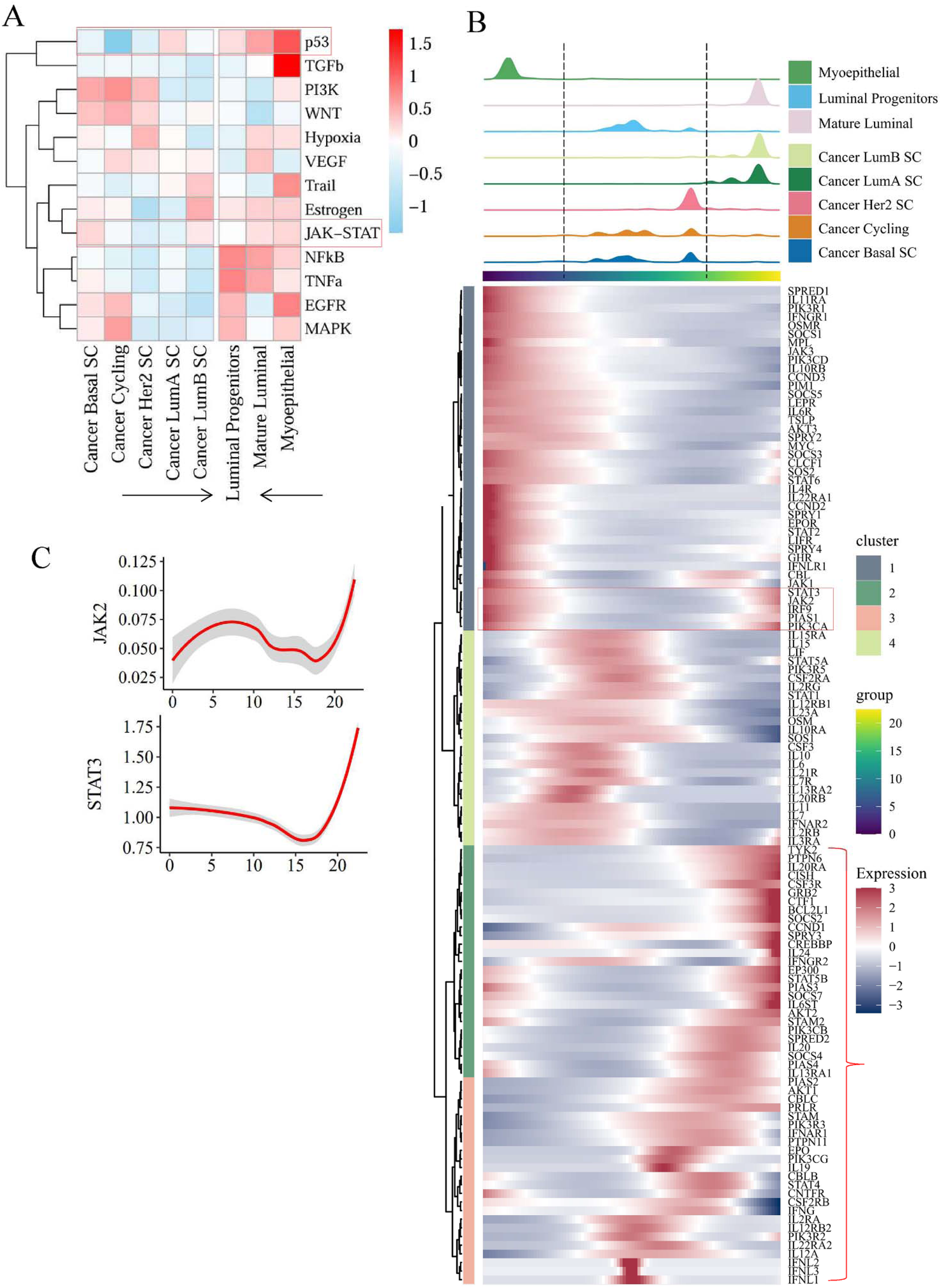
Progeny pathway activity and Monocle2 profiling. (A) Progeny pathway activity of epithelial cells. (B) Pseudotime trajectory analysis in epithelial cells: heatmap of JSP gene expression. Red brackets: T cell differentiation-specific genes. (C) Relationship between pseudotime score and gene expression (JAK2, STAT3).

**Figure S3.**
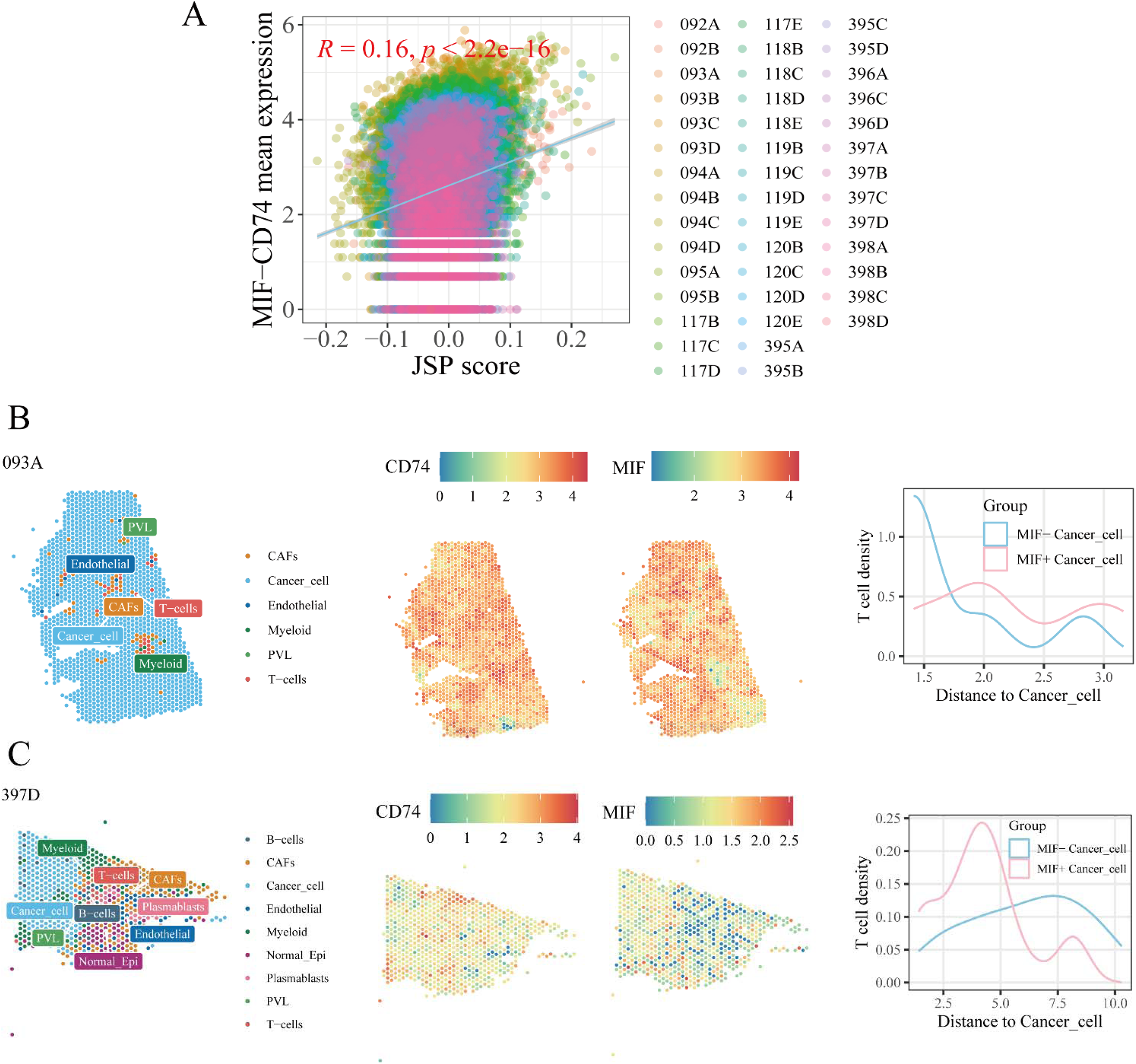
Spatial transcriptomic analysis validates the MIF□CD74 axis. (A) Correlation between the mean expression of MIF□CD74 and JSP score. (B□C) Representative cases 093A (B) and 397D (C) from the GSE210616 dataset. Left panels: cell type annotation; middle panels: spatial expression of MIF and CD74; right panels: spatial neighborhood analysis of MIF⁺ and MIF⁻ tumor cells.

**Figure S4.**
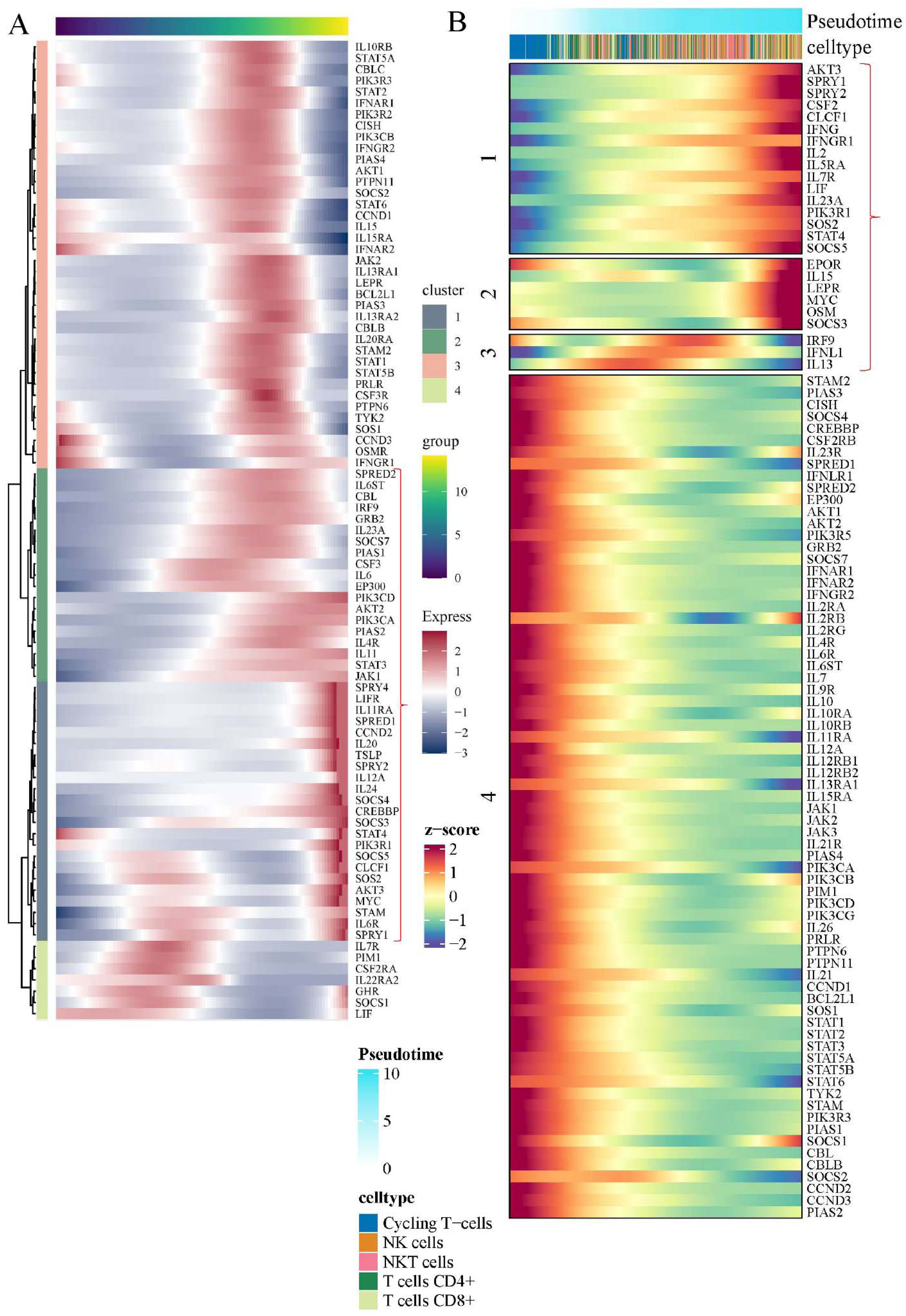
Pseudotime trajectory analysis of the JAK-STAT pathway in normal epithelial cells and T cells. Differentiation trajectory of normal epithelial cells (A) and T cells (B). Genes associated with JSP that exhibited elevated expression during late differentiation were designated as cell differentiation-dependent genes, marked with red brackets.

**Figure S5.**
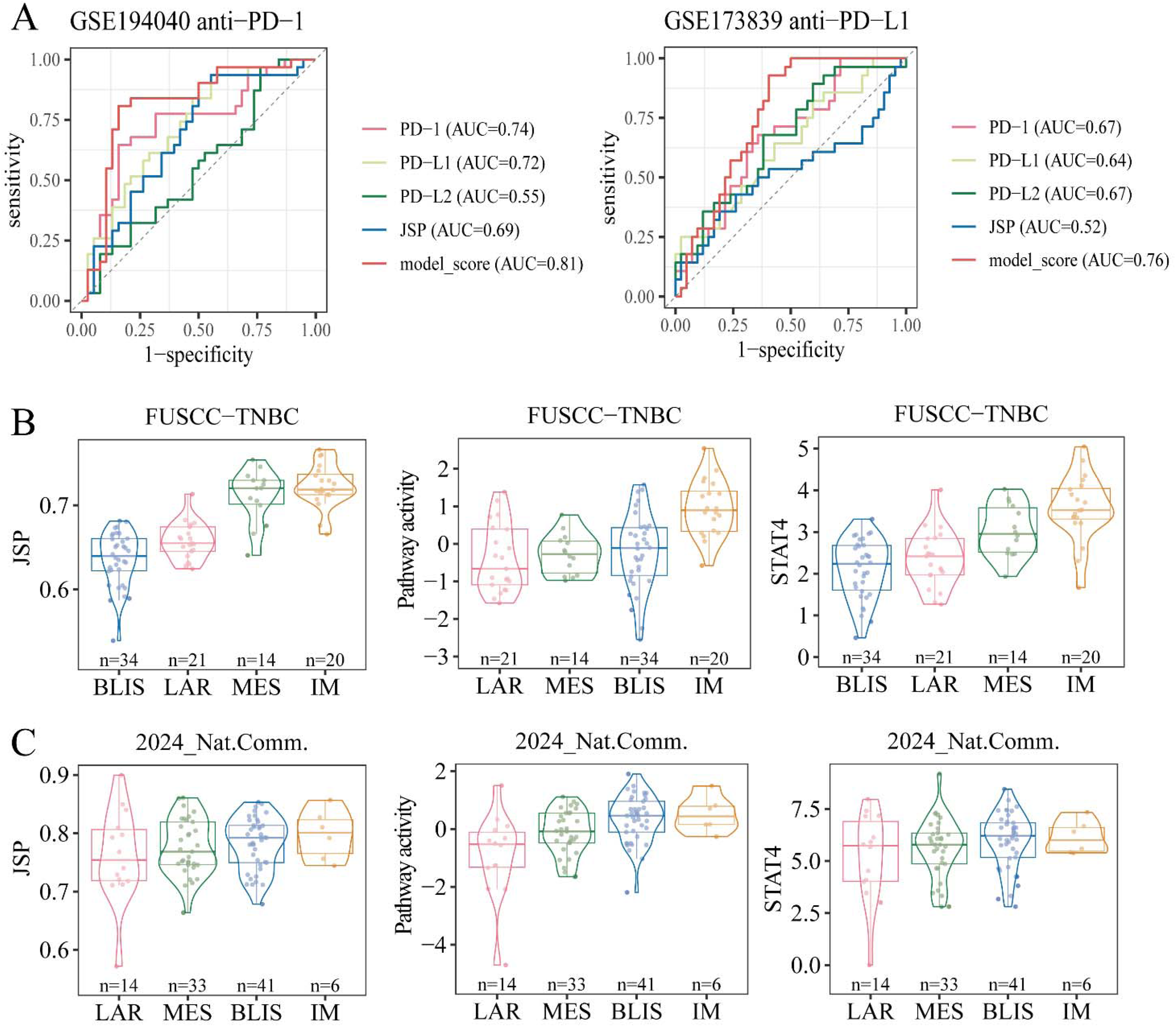
Combined predictive model and heterogeneity of the JAK-STAT pathway in TNBC Subtypes. (A) ROC curves of PDCD1(PD-1), CD274(PD-L1), PDCD1LG2(PD-L2), JSP score and combined model score. (B) FUSCC-TNBC cohort. (C) 2024_Nat.Comm. cohort. Left panels: JSP scores; middle panels: JAK-STAT pathway activity; right panels: STAT4 expression. LAR: luminal androgen receptor, IM: immunomodulatory, BLIS: basal□like immune□suppressed, MES: mesenchymal□like.

**Figure S6.**
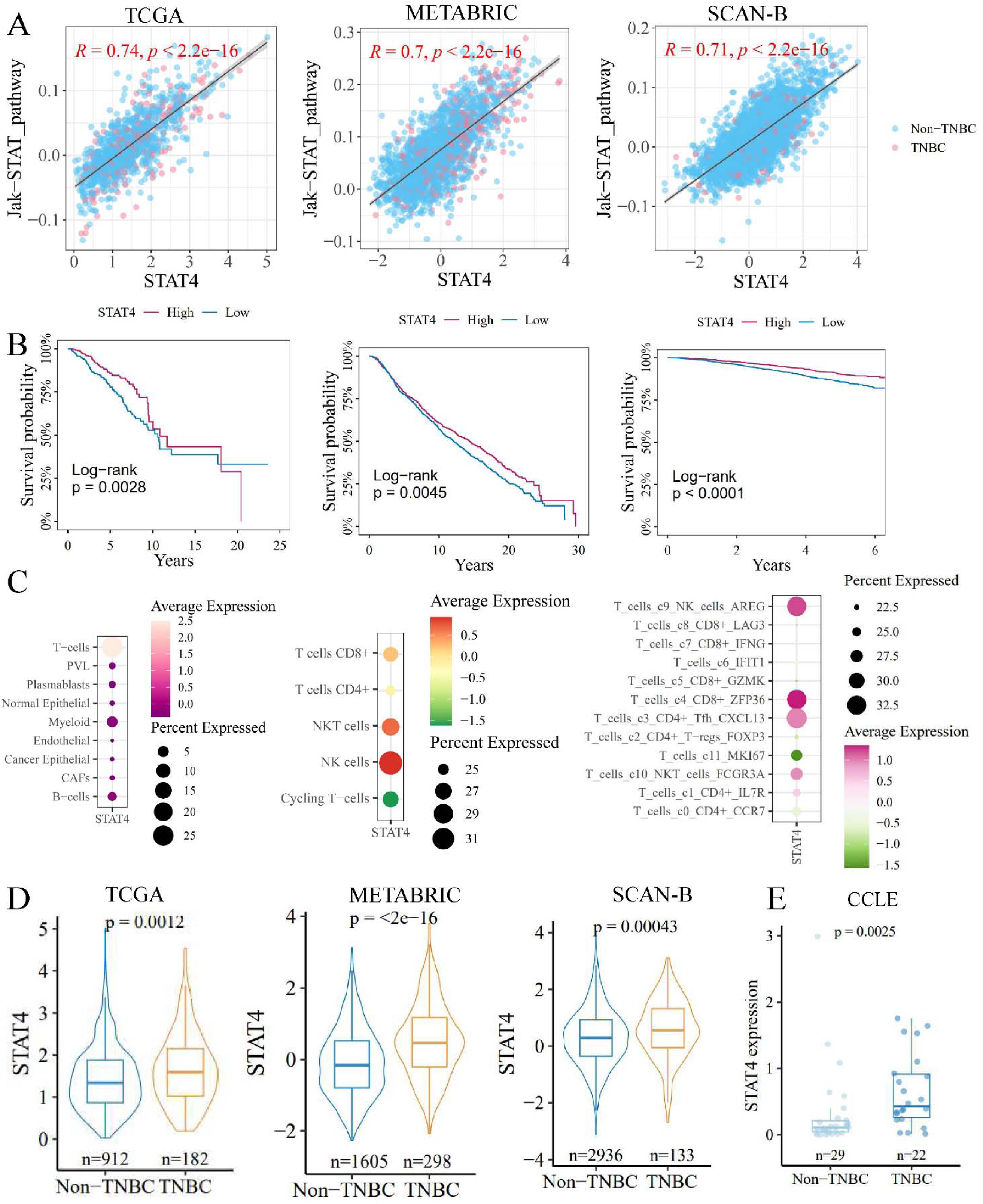
STAT4 as a representative of the JAK-STAT pathway. (A) Correlation between STAT4 expression and the JAK-STAT pathway score across the TCGA, METABRIC, and SCAN-B cohorts. (B) Survival analysis of STAT4 within these cohorts, using the median as the threshold. (C) Expression patterns of STAT4 in a breast cancer single-cell dataset (GSE176078), highlighting major subtypes, T-cell minor subtypes, and T-cell subsets. (D) Variation in STAT4 expression between TNBC and non-TNBC samples. (E) Expression of STAT4 in breast cancer cell lines, categorized into TNBC and non-TNBC subgroups.

**Figure S7.**
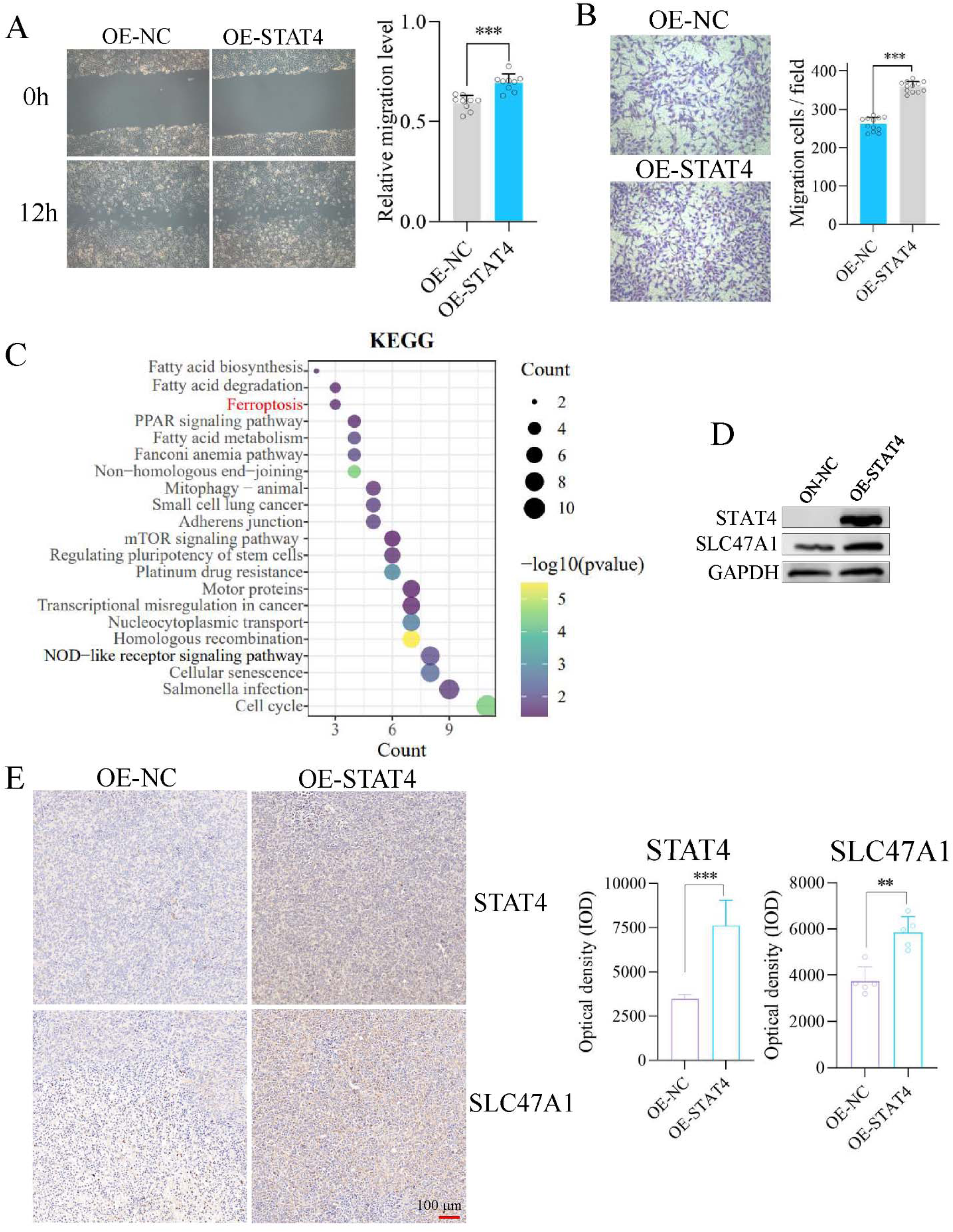
STAT4 up-regulates SLC47A1 to involve ferroptosis in breast cancer cells. (A-B) Wound healing (A) and cell migration assays (B) were performed following STAT4 overexpression in MDA-MB-231 cells. (C) KEGG pathway enrichment analysis of differentially expressed genes following STAT4 overexpression. (D) Upregulation of SLC47A1 upon STAT4 overexpression. (E) Expression levels of STAT4 and SLC47A1 in xenograft tumors in nude mice. Significance: *, *p* < 0.05; ** *p* < 0.01; ***, *p* < 0.001.

**Figure S8.**
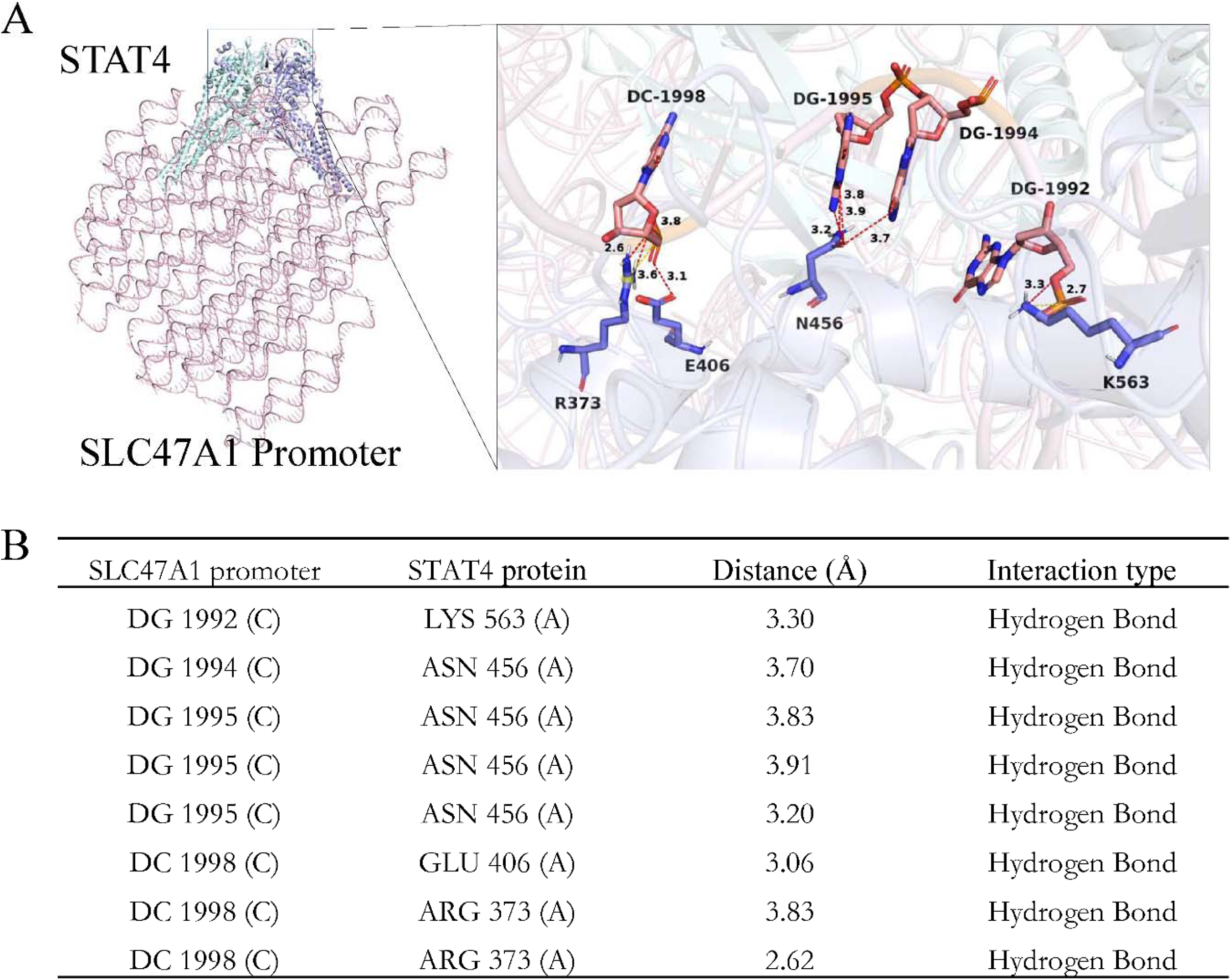
STAT4 binds to the SLC47A1 promoter. (A) Hydrogen bond interactions between STAT4 and its binding sites on the SLC47A1 promoter, with a binding energy of −46.28 kcal/mol. (B) Positions and corresponding distances of these hydrogen bonds.

