## Supplementary_Table for "JAK-STAT Pathway Heterogeneity Governs Immunotherapy Response in Breast Cancer"

**Supplementary Table 1** JAK-STAT pathway genes

AKT3

SPRY3

SPRY1

SPRY2

STAM2

IRF9

PIAS3

IL24

CISH

IL22RA2

SOCS4

CNTF

CNTFR

CREBBP

CSF2

CSF2RA

CSF2RB

CSF3

CSF3R

CSH1

CTF1

IL23R

SPRED1

IFNLR1

SPRED2

EP300

EPO

EPOR

AKT1

AKT2

CLCF1

PIK3R5

CBLC

GH1

GH2

GHR

IFNL2

IFNL3

IFNL1

GRB2

IL19

SOCS7

IFNE

IFNA1

IFNA2

IFNA4

IFNA5

IFNA6

IFNA7

IFNA8

IFNA10

IFNA13

IFNA14

IFNA16

IFNA17

IFNA21

IFNAR1

IFNAR2

IFNB1

IFNG

IFNGR1

IFNGR2

IFNW1

IL2

IL2RA

IL2RB

IL2RG

IL3

IL3RA

IL4

IL4R

IL5

IL5RA

IL6

IL6R

IL6ST

IL7

IL7R

IL9

IL9R

IL10

IL10RA

IL10RB

IL11

IL11RA

IL12A

IL12B

IL12RB1

IL12RB2

IL13

IL13RA1

IL13RA2

IL15

IL15RA

JAK1

JAK2

JAK3

LEP

LEPR

LIF

LIFR

MPL

MYC

OSM

IL20

IL21R

IL22

IL23A

PIAS4

PIK3CA

PIK3CB

PIM1

PIK3CD

PIK3CG

PIK3R1

PIK3R2

IL20RA

IL20RB

IL26

PRL

PRLR

IFNK

PTPN6

PTPN11

IL22RA1

IL21

CCND1

BCL2L1

CRLF2

SOS1

SOS2

STAT1

STAT2

STAT3

STAT4

STAT5A

STAT5B

STAT6

TPO

TYK2

STAM

SPRY4

PIK3R3

TSLP

PIAS1

SOCS1

CBL

CBLB

SOCS2

CCND2

CCND3

SOCS3

PIAS2

OSMR

SOCS5

**Supplementary Table 2** The genes for venn plot.

| **Tumor** | **Epi** | **Tcell** |
| --- | --- | --- |
| IFNL1 | SPRY1 | AKT3 |
| IFNL3 | IL6R | SPRY1 |
| IFNL2 | STAM | SPRY2 |
| IL12A | MYC | CSF2 |
| IL22RA2 | AKT3 | CLCF1 |
| PIK3R2 | SOS2 | IFNG |
| IL12RB2 | CLCF1 | IFNGR1 |
| IL2RA | SOCS5 | IL2 |
| IFNG | PIK3R1 | IL5RA |
| CSF2RB | STAT4 | IL7R |
| CNTFR | SOCS3 | LIF |
| STAT4 | CREBBP | IL23A |
| CBLB | SOCS4 | PIK3R1 |
| IL19 | IL24 | SOS2 |
| PIK3CG | IL12A | STAT4 |
| EPO | SPRY2 | SOCS5 |
| PTPN11 | TSLP | EPOR |
| IFNAR1 | IL20 | IL15 |
| PIK3R3 | CCND2 | LEPR |
| STAM | SPRED1 | MYC |
| PRLR | IL11RA | OSM |
| CBLC | LIFR | SOCS3 |
| AKT1 | SPRY4 | IRF9 |
| PIAS2 | JAK1 | IFNL1 |
| IL13RA1 | STAT3 | IL13 |
| PIAS4 | IL11 |  |
| SOCS4 | IL4R |  |
| IL20 | PIAS2 |  |
| SPRED2 | PIK3CA |  |
| PIK3CB | AKT2 |  |
| STAM2 | PIK3CD |  |
| AKT2 | EP300 |  |
| IL6ST | IL6 |  |
| SOCS7 | CSF3 |  |
| PIAS3 | PIAS1 |  |
| STAT5B | SOCS7 |  |
| EP300 | IL23A |  |
| IFNGR2 | GRB2 |  |
| IL24 | IRF9 |  |
| CREBBP | CBL |  |
| SPRY3 | IL6ST |  |
| CCND1 | SPRED2 |  |
| SOCS2 |  |  |
| BCL2L1 |  |  |
| CTF1 |  |  |
| GRB2 |  |  |
| CSF3R |  |  |
| CISH |  |  |
| IL20RA |  |  |
| PTPN6 |  |  |
| TYK2 |  |  |
